# Macromolecular Crowding Tailors the Microtubule Cytoskeleton Through Tubulin Modifications and Microtubule-Associated Proteins

**DOI:** 10.1101/2023.06.14.544846

**Authors:** Yusheng Shen, Kassandra M. Ori-McKenney

## Abstract

Cells remodel their cytoskeletal networks to adapt to their environment. Here, we analyze the mechanisms utilized by the cell to tailor its microtubule landscape in response to changes in osmolarity that alter macromolecular crowding. By integrating live cell imaging, *ex vivo* enzymatic assays, and *in vitro* reconstitution, we probe the impact of acute perturbations in cytoplasmic density on microtubule-associated proteins (MAPs) and tubulin posttranslational modifications (PTMs), unraveling the molecular underpinnings of cellular adaptation via the microtubule cytoskeleton. We find that cells respond to fluctuations in cytoplasmic density by modulating microtubule acetylation, detyrosination, or MAP7 association, without differentially affecting polyglutamylation, tyrosination, or MAP4 association. These MAP-PTM combinations alter intracellular cargo transport, enabling the cell to respond to osmotic challenges. We further dissect the molecular mechanisms governing tubulin PTM specification, and find that MAP7 promotes acetylation by biasing the conformation of the microtubule lattice, and directly inhibits detyrosination. Acetylation and detyrosination can therefore be decoupled and utilized for distinct cellular purposes. Our data reveal that the MAP code dictates the tubulin code, resulting in remodeling of the microtubule cytoskeleton and alteration of intracellular transport as an integrated mechanism of cellular adaptation.

## Introduction

Adaptation is key for survival. During their lifetimes, cells will be exposed to a variety of conditions to which they must adapt to maintain homeostasis. A part of this adaptation response is an expansive repertoire of proteins to remodel their cytoskeletal networks. The microtubule cytoskeleton is dynamically regulated by a range of microtubule-associated proteins (MAPs), tubulin posttranslational modifications (PTMs), and the expression of distinct tubulin isotypes [1–5]. Structural MAPs, such as MAP7, MAP4, and tau, that co-purify with polymerized brain tubulin and bind along the shaft of the lattice, can stabilize and bundle microtubules, alter the spacing of tubulin dimers within the lattice, and facilitate or inhibit kinesin and dynein motor transport [6–12]. Tubulin PTMs include luminal acetylation within the unstructured *α*K40-loop, and complex chemical modifications of the cytosolic unstructured C-terminal tails including tyrosination/detyrosination and polyglutamylation [1, 2]. Tubulin PTMs can regulate the dynamicity and stability of microtubules, alter the biophysical parameters of the microtubule lattice, and recruit specific effectors [13–17]. These two systems of regulation are referred to as the “MAP code” and the “tubulin code” [2–4, 9, 18–20]. We are only beginning to understand the interplay between these two codes. Some MAPs that have been shown to control microtubule dynamics, such as CLIP170, or motor onloading, such as dynactin, are affected by tyrosination status [16, 21, 22], but less is known about the effects of these PTMs on structural MAPs. Recent work has shown that doublecortin is necessary for the maintenance of polyglutamylation in neurons [23], which directly promotes the binding of tau to microtubules [24]. How other MAPs and PTMs communicate and how cells utilize combinations of MAPs and PTMs for specific purposes are open questions.

Dynamic microtubules exist in a dense cytoplasmic milieu. The mechanical properties of cytoplasm influence several cellular functions; however, cells can experience variations in osmolarity, and therefore macromolecular crowding, in response to metabolic, oxidative, or inflammatory stress and as a result of aging or tumorigenesis [25–29]. Therefore, the cell must develop strategies to respond to these challenges. Prior studies have shown that increases in viscosity or macromolecular crowding facilitate microtubule stability, suppressing both microtubule polymerization and depolymerization, as well as reduce the efficiency and speed of motor protein transport [30–33]. We have little understanding of how cells respond to these global effects at the microtubule level through MAPs and tubulin PTMs, or how the cell remodels its microtubule network to adapt to a changing environment. Unraveling these mysteries holds the potential to uncover novel mechanisms underlying cellular adaptability and pathological transformation.

Here we use osmotic challenges to acutely manipulate the macromolecular crowding of cytoplasm and investigate how the cell integrates the MAP and tubulin codes to tailor its microtubule cytoskeleton in response to these changes. We reveal that increasing cytoplasmic density promotes MAP7 association with microtubules and enhances microtubule acetylation. In contrast, decreasing cytoplasmic density reduces the association of MAP7 with microtubules and augments microtubule detyrosination. Acetylation and detyrosination of microtubules are usually tightly linked in cells, but surprisingly, we find that the presence or absence of MAP7 selects for one modification over the other, thus decoupling them to be used in distinct cellular contexts. Using *ex vivo* acetylation assays and *in vitro* reconstitution experiments, we show that MAP7 binding promotes acetylation by allosterically altering the lattice to facilitate *α*-tubulin acetyl transferase-1 (*α*TAT1) activity and prevents detyrosination by sterically interfering with vasohibin 1 (VASH1-SVBP) binding. Finally, we perform live cell imaging to reveal that increasing macromolecular crowding does not hinder all microtubule-based transport, but selects against retrograde transport of early endosomes without impeding anterograde transport of secretory vesicles. In contrast, decreased crowding results in the opposite effect. Altering cytoplasmic density therefore selects for specific modes of transport not simply due to changes in the physical properties of the cytoplasm, but through alteration of MAP association which directly impacts tubulin PTMs. Therefore, we reveal that the tubulin code directly depends on the MAP code. Our results provide insight into how the microtubule cytoskeleton can be remodeled by distinct MAPs and PTMs to efficiently respond to variations in macromolecular crowding.

## Results

### Cytoplasm density differentially tunes tubulin PTMs

We sought to understand how cells modulate their microtubule cytoskeletons in response to acute alterations in cytoplasmic density. To manipulate cytoplasmic density, we exposed cells to extracellular osmotic environments ranging from hypoosmotic to hyperosmotic conditions (200–400 mOsm), in order to either increase or decrease cell volume through the intake or loss of water, respectively [30, 34, 35]. We produced specific osmolarity conditions by varying the amount of mannitol, a cell-membrane-impermeable non-toxic sugar, to a hypotonic base solution while maintaining a fixed ionic strength [36, 37]. To directly measure effects of these osmotic challenges on cytoplasmic density, we expressed 40 nm diameter genetically encoded multimeric (GEM) nanoparticles in BEAS-2B cells, a human lung epithelial line (Fig. 1A). BEAS-2B cells typically spread as a thin lamella when cultured on glass, largely confining the diffusion of GEMs in 2 dimensions, which we imaged using total internal reflection fluorescence (TIRF) microscopy (Fig. 1A). Using single-particle tracking [38], we tracked GEM diffusion and measured their mean-squared displacements (MSDs) for cells that were acutely treated with either isotonic (310 mOsm, control), hypotonic (250 mOsm) and hypertonic (400 mOsm) solutions (Fig. 1B). As has been observed for other cell types [39], the measured MSDs were linear functions of delay time *τ*, with measured diffusion coefficients of 0.068±0.012, 0.19±0.05, and 0.039±0.005 *μm*^2^*/s* for isotonic, hypotonic, and hypertonic osmolarities, respectively. The changes of GEM diffusion coefficients indicate that cytoplasm density is indeed increased upon hypotonic treatment and decreased upon hypertonic treatment. We conclude that acute osmotic challenges can be used to manipulate the macromolecular crowding of the cytoplasm in living cells.

**FIG. 1.**
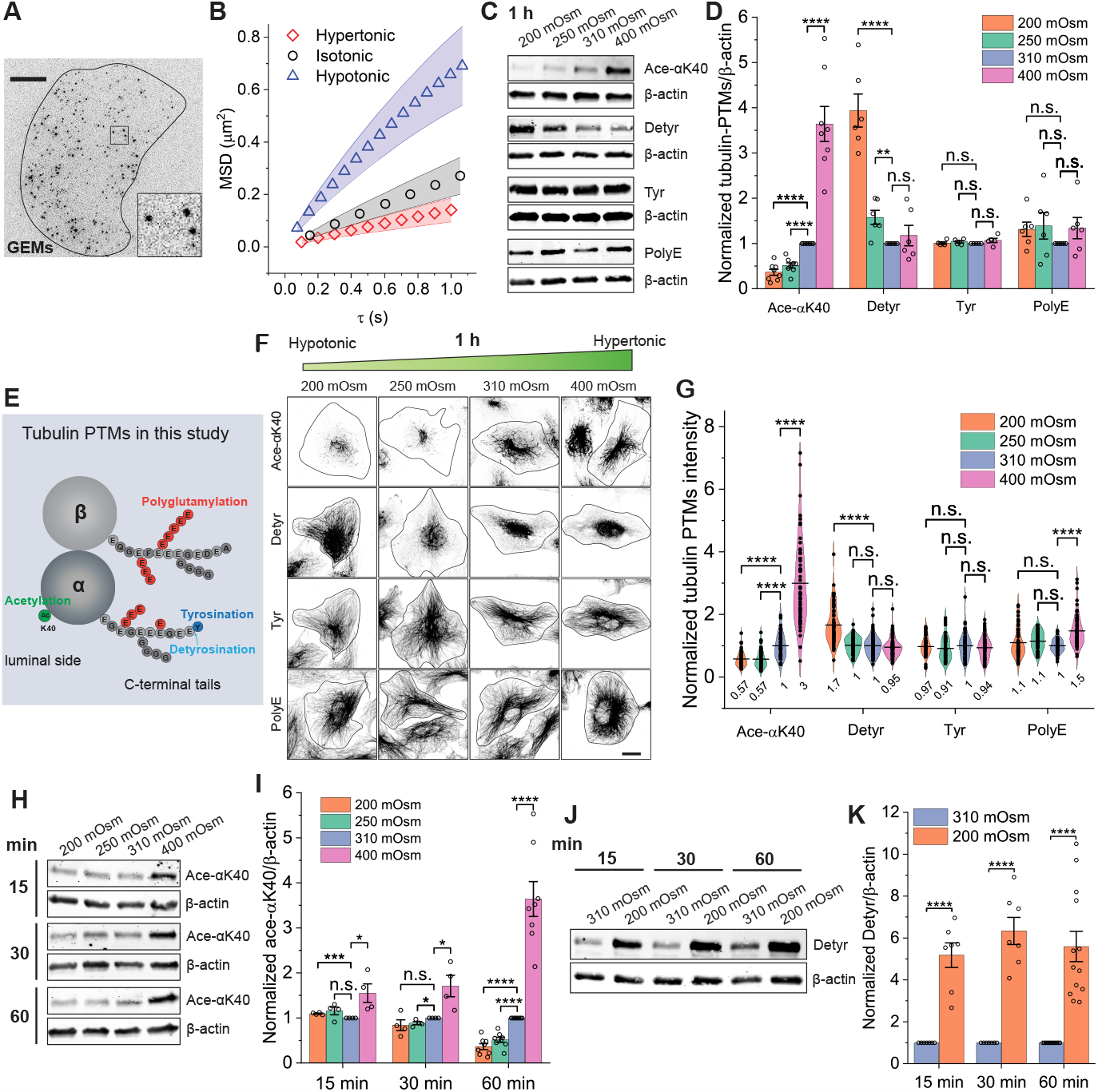
Cytoplasm crowdedness differentially tunes tubulin post-translational modifications. (A) Image of a BEAS-2B cell expressing 40 nm-GEMs visualized by TIRF microscopy. Inset: magnified view of GEMs. Scale bar: 10 *μ*m. (B) Averaged mean square displacement (MSD) of GEMs as a function of delay time, *τ*, for cells treated with either isotonic (310 mOsm, black circles), hypotonic (250 mOsm, blue triangles) or hypertonic (400 mOsm, red diamonds) solutions. Cells were imaged between 5 to 65 min incubation with corresponding solutions. Shaded areas indicate S.D. The total trajectories analyzed are n = 13689, 26392 and 5569, and n = 13, 10 and 10 cells from 2 independent experiments. (C and D) Western-blots (WB) and corresponding quantification showing relative changes of tubulin PTMs in response to different osmotic treatments for 1 hr. Acetylation at Lys40 (Ace-*α*K40), detyrosination (Detyr), tyrosination (Tyr), and polyglutamylation (PolyE) were assessed relative to the loading control, *β*-actin. Ace-*α*K40, n = 8, \*\*\*\**p <* 0.0001; Detyr, n = 6, \*\*\*\**p <* 0.0001, \*\**p* = 0.0041; Tyr, n = 5; PolyE, n = 6. n.s. indicates *p >* 0.05. (E) Relative positions of tubulin PTMs to the globular, highly structured tubulin bodies and the unstructured C-terminal tails. (F and G) Immunostaining analysis showing cellular distribution and relative changes of tubulin PTMs in response to different osmotic treatments for 1 hr. Black lines indicate the cell edge. Scale bar: 20 *μ*m. Ace-*α*K40, n = 50, 53, 51 and 58 cells from 2 independent experiments, \*\*\*\**p <* 0.0001; Detyr, n = 69, 77, 77 and 86 cells from 3 independent experiments, \*\*\*\**p <* 0.0001; Tyr, n = 45, 54, 53 and 59 cells from 2 independent experiments; PolyE, n = 61, 59, 62 and 63 cells from 2 independent experiments, \*\*\*\**p <* 0.0001. (H and I) WBs and corresponding quantification showing the temporal changes of cellular ace-*α*K40 following osmotic challenges. 15 min: n = 4, \*\*\**p* = 0.00021, \**p* = 0.036; 30 min: n = 4, \**p* = 0.015 and 0.025; 60 min: n = 8, \*\*\*\**p <* 0.0001. (J and K) WBs and corresponding quantification showing the temporal changes of cellular Detyr following iso- and hypo-osmotic challenges. 15 min: n = 7, \*\*\*\**p <* 0.0001; 30 min: n = 7, \*\*\*\**p <* 0.0001; 60 min: n = 13, \*\*\*\**p <* 0.0001. All graphs display all datapoints with means. Error bars indicate s.e.m., unless otherwise noted. All *p*-values were calculated using a student’s t-test.

We first examined how tubulin PTMs respond to variations in cytoplasm density. We subjected BEAS-2B cells to one-hour osmotic challenges and assessed the relative effects on tubulin PTMs using both immunoblotting and immunostaining, which reveal average cell population responses and changes at the single cell level, respectively. We analyzed the *α*-tubulin luminal modification, acetylation of lysine 40 (“acetylation” hereafter), the *α*-tubulin C-terminal tail modifications, detyrosination (Detyr) and tyrosination (Tyr), and the *α*- and *β*-tubulin C-terminus modification, polyglutamylation (PolyE) (Fig. 1C-G) [2]. Unexpectedly, acetylation and detyrosination, which are both considered markers for stabilized microtubules and often occur on the same subsets of microtubules within cells [2, 13], responded differently to osmotic shifts. As measured by immunoblotting, increasing cytoplasmic density caused a 3.5-fold increase in *α*-tubulin acetylation levels, whereas decreasing the cytoplasmic density resulted in a 64% decrease in *α*-tubulin acetylation levels compared to the isotonic control (Fig. 1C-D). Interestingly, decreasing cytoplasmic density resulted in increased detyrosination levels (1.6 and 3.9-fold increase for 250 mOsm and 200 mOsm, respectively), whereas *α*-tubulin tyrosination and polyglutamylation levels were insensitive to these osmotic shifts (Fig. 1C-D). Similar results were observed by immunostaining (Fig. 1F-G). Again, hypertonic treatment caused a 3-fold increase in acetylation staining (Fig. 1F-G). Upon hypotonic treatment, acetylation staining was greatly diminished, whereas detyrosination levels were greatly increased and expanded within cells compared to the isotonic control. Both tyrosination and polyglutamylation signals homogeneously decorated the microtubules and remained relatively unchanged for all the conditions (Fig. 1F-G); however, we did observe a 1.5-fold increase in polyglutamylation in the hyperosmotic condition (Fig. 1F-G). The immunoblotting and immunostaining results were broadly consistent, demonstrating that cytoplasm density differentially tunes tubulin PTMs. Furthermore, we observed similar differential regulation of tubulin PTMs in HeLa and RPE-1 cells (Fig. S1A-B), indicating a conserved response in various types of epithelial cells. Notably, changes in detyrosination levels were far more pronounced in the cancerous epithelial HeLa cell line, which showed a 40% decrease in response to the hyperosmotic condition and a 4.2-fold increase in the hypoosmotic condition (Fig. S1), indicating that detyrosination levels in cancer cells may be more sensitive to changes in cytoplasm crowdedness. Thus, our results reveal that the two most prominent markers of stable microtubules, acetylation and detryosination, are separable under specific cellular conditions.

We observed the most pronounced effects on acetylation upon increasing cytoplasmic density, and on detyrosination upon decreasing density. We were therefore curious about the temporal response of these two modifications considering acetylation occurs within the lumen of the microtubule, while detyrosination occurs at the more accessible C-terminal tail [40–43]. Upon hypertonic treatment, acetylation levels gradually increased (Fig. 1H-I, 1.5, 1.7 and 3.6-fold for 15, 30 and 60 min, respectively), whereas detyrosination levels rose sharply after only 15 min of hypotonic treatment (Fig. 1J-K), supporting the notion that it may take longer for acetylation enzymes to access the microtubule lumen [44]. In hypoosmotic conditions, the decrease of acetylation becomes evident only after 1 hr (Fig. 1H-I), suggesting that cells need more time to process the removal rather than the addition of this modification.

### Cytoplasm density has distinct effects on MAPs

Tubulin PTMs have been shown to impact the association of microtubule-associated proteins (MAPs) with microtubules [1–3, 21, 45]. Because our previous results revealed that changes in cytoplasmic density alter the microtubule PTM landscape within cells, we wondered if these changes might also influence the interactions of MAPs with microtubules. To test this, we performed immunostaining to localize the subcellular distribution of four prominent non-neuronal MAPs in response to varying osmotic conditions [46–50]. We first confirmed the expression of MAP7 and MAP4 in BEAS-2B cells, while MAP9 was only present in cells undergoing mitosis, and DCLK1 expression was not detected (data not shown). We therefore examined the subcellular distribution and relative expression of endogenous MAP7 and MAP4 in response to osmotic challenges (Fig. 2A-C). Upon hypotonic treatment, MAP7 dissociated from microtubules and became diffusive within the cytosol, especially in the lowest osmolarity condition, whereas MAP4 localization along microtubules appeared unchanged on microtubules (Fig. 2A). Compared to the isotonic control, MAP7 intensity increased 1.2-fold in the hyperosmotic condition, but decreased in both hypoosmotic conditions (37% and 19% for 200 and 250 mOsm, respectively). In contrast, MAP4 intensity on microtubules increased under all conditions examined (Fig. 2B-C). We next used immunoblotting to assess overall changes in MAP7 or MAP4 protein levels. Intriguingly, MAP7 was modestly upregulated upon hypotonic treatment and downregulated upon hypertonic treatment, whereas MAP4 levels remained unchanged (Fig. 2D-E). The observed effects on MAP7 protein levels might be a cellular response to alterations in the association of MAP7 with microtubules (Fig. 2A). Osmotic regulation of MAP7 levels was also observed in RPE-1 cells, but not in HeLa cells, where MAP7 levels are physiologically higher (Fig. S2). To determine the effects on MAP7 and MAP4 microtubule binding behaviors under different osmotic treatments, we used a truncated CMV promoter to transiently express EGFP-tagged MAPs at reduced levels, allowing us to monitor MAP dynamics in live cells [51]. EGFP-MAP7 was able to associate with microtubules in the isoosmotic and hyperosmotic conditions, but did not bind microtubules in the hypoosmotic condition, whereas EGFPMAP4 bound to microtubules in all conditions (Fig. 2F). These results demonstrate that MAP7 and MAP4 respond differently to the changes in cytoplasm density. Unlike MAP4, MAP7 association with microtubules is differentially affected by osmotic conditions, which interestingly correlates with the changes in acetylation levels we previously observed (Fig. 1).

**FIG. 2.**
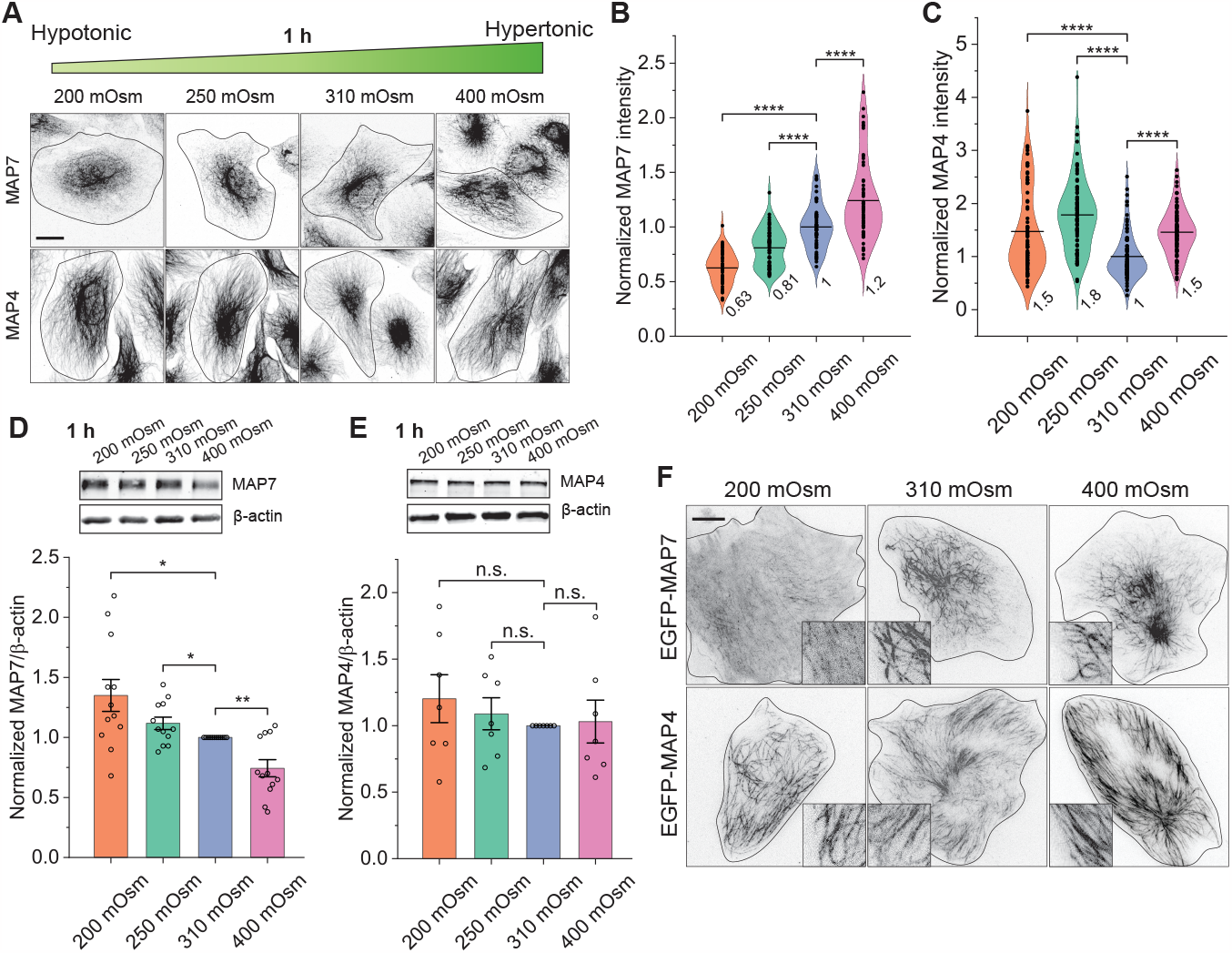
Cytoplasm crowdedness differentially tunes microtubule-associated proteins. (A-C) Immunostaining analysis showing the cellular distribution and relative intensity changes of endogenous MAP7 and MAP4 in response to osmotic treatments for 1 hr. Black lines indicate cell edge. Scale bar: 20 *μ*m. MAP7: n = 50, 53, 51 and 58 cells from 2 independent experiments, \*\*\*\**p <* 0.0001; MAP4: n = 78, 82, 83 and 80 cells from 2 independent experiments, \*\*\*\**p <* 0.0001. (D and E) WBs and corresponding quantification showing the relative changes of MAP7 and MAP4 in response to osmotic treatments for 1 hr. *β*-actin, loading control. MAP7: n = 12, \**p* = 0.015 and 0.033, \*\**p* = 0.0017; MAP4: n = 7, no significant changes. (F) Images of BEAS-2B cells transiently expressing EGFP-MAP7 and EGFP-MAP4 showing the distribution of MAPs following osmotic treatments for 1 hr. A truncated CMV (DeltaCMV) promoter was used to express MAPs at reduced levels [51]. Insets show a magnified portion of each image. Scale bar: 10 *μ*m. All graphs display all data points with means. Error bars indicate s.e.m. All *p*-values were calculated using a student’s t-test.

### MAP7 promotes and protects microtubule acetylation

In BEAS-2B cells, MAP4 homogenously decorated microtubules, whereas MAP7 largely localized to the perinuclear region (Fig. 3A). Acetylated and detyrosinated microtubules are also concentrated at the perinuclear region, but with distinct spatial patterns, in agreement with a previous finding that acetylation and detyrosination decorate distinct microtubule regions [52, 53]. Interestingly, acetylation largely colocalized with MAP7 (Fig. 3A). Combined with our above results, this suggests there may be an interplay between tubulin acetylation and MAP7 association. To define this interplay, we first examined the effect on acetylation levels upon exogenous expression of MAP7. We transiently expressed full length MAP7 (EGFP-MAP7) or a C-terminal fragment of MAP7 (EGFP-MAP7-Cter) that lacks the microtubule-binding domains as a control. We found that expression of full-length MAP7 caused a 1.9-fold increase of acetylation compared to MAP7-Cter (Fig. 3BC). Furthermore, acetylation and MAP7 signals largely overlapped along microtubules (Fig. 3B). We next examined the MAP7 response to increased acetylation levels by treating the cells for 1 hr with tubacin, a drug that inhibits histone deacetylase 6 (HDAC6) [54]. Both the cellular distribution and levels of MAP7 protein remained unchanged compared to controls, despite the *∼*9-fold increase in acetylation signal that expanded to encompass nearly all microtubules (Fig. 3D-F). These results are consistent with a previous finding that loss of MAP7 reduces acetylation in nascent branches of neurons [10], and suggests that MAP7 may promote acetylation in BEAS2B cells, but not *vice versa*.

**FIG. 3.**
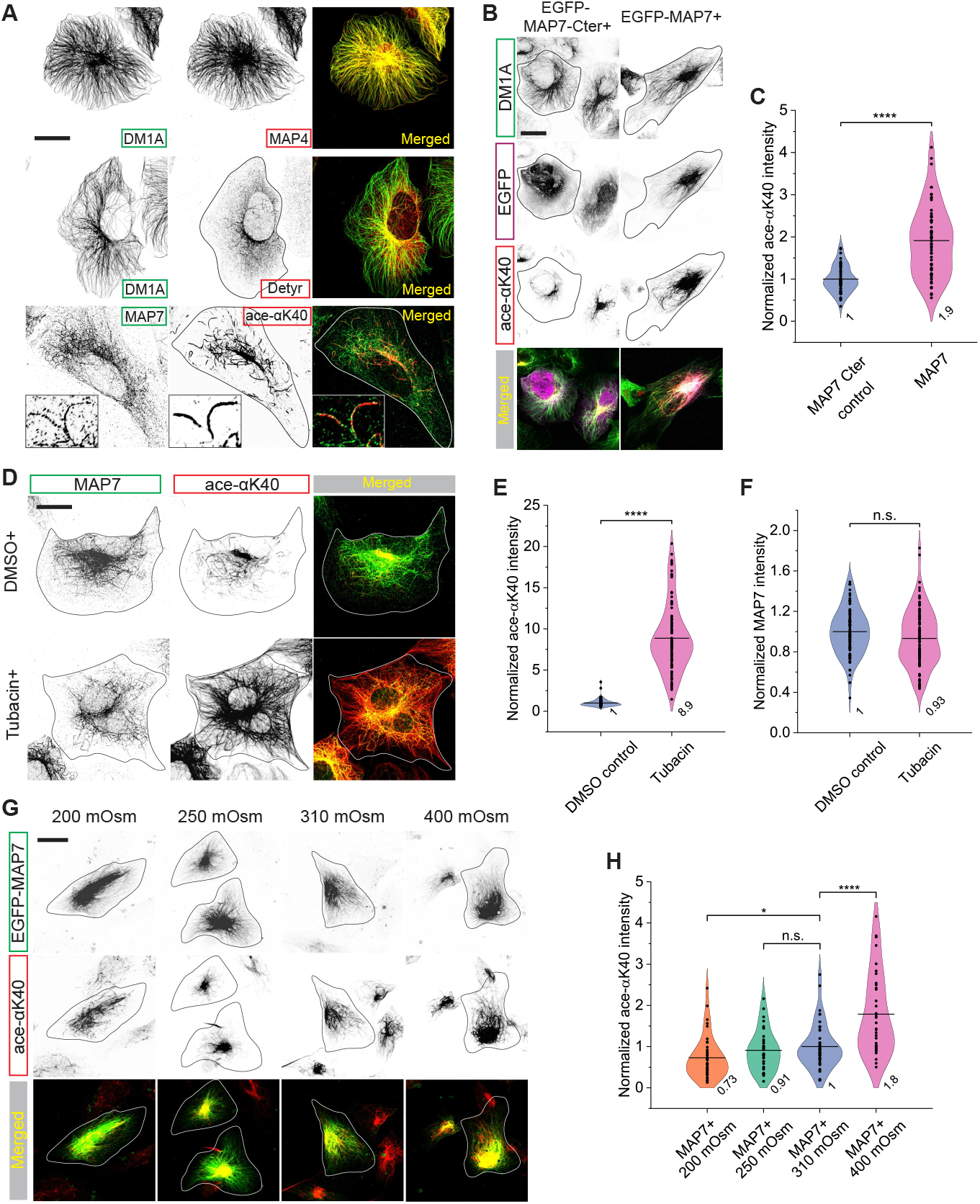
MAP7 promotes and protects microtubule acetylation. (A) Immunostaining analysis showing the relative cellular distribution of MAP4, MAP7, Detyr, and Ace-*α*K40 compared to total *α*-tubulin (DM1A). Acetylation decorates microtubules continuously as patches, whereas detyrosination mostly appears as diffusive puncta. Insets show a magnified portion of the images in which Ace-*α*K40 is highly colocalized with MAP7. (B and C) Representative images and quantification comparing the fluorescence intensity of Ace-*α*K40 for cells transiently expressing either the C-terminal region of MAP7 (EGFP-MAP7-Cter, control) or full length MAP7 (EGFP-MAP7). n = 49 and 61 cells, respectively, from 2 independent experiments, \*\*\*\**p <* 0.0001. (D-F) Representative images and quantification comparing the fluorescence intensity of Ace-*α*K40 and MAP7 for cells that were either treated with DMSO (control) or tubacin (an inhibitor of HDAC6, 5 *μ*M) for 1 hr. n = 95 and 100 cells, respectively, from 3 independent experiments, \*\*\*\**p <* 0.0001. (G and H) Representative images and quantification comparing the fluorescence intensity of Ace-*α*K40 for cells transiently expressing MAP7 in response to different osmotic challenges for 1 hr. n = 40, 32, 43 and 35 cells from 2 independent experiments, \**p* = 0.023, \*\*\*\**p <* 0.0001. The acetylation level was significantly higher than that without MAP7 expression for the 250 mOsm condition (\*\*\**p* = 0.00011, Fig. 1G). Scale bars: 20 *μ*m. All graphs display all data points with means. All *p*-values were calculated using a student’s t-test.

We next asked how exogenous expression of MAP7 would affect microtubule acetylation under different osmotic conditions, considering hypoosmotic treatment results in a decrease in both acetylation and MAP7 association (Figs. 1F-G and 2A-B). We subjected EGFPMAP7 expressing cells to 1 hr osmotic treatments, then measured the fluorescent intensity of microtubule acetylation by immunostaining (Fig. 3G-H). Upon expression of MAP7, acetylation levels were significantly higher in the 250 mOsm condition (Fig. 1G and Fig. 3H), suggesting that MAP7 promoted microtubule acetylation under osmotic conditions that normally resulted in a strong reduction of this modification. We conclude that MAP7 binding along microtubules may protect acetylated microtubules from deacetylation.

### Cytoplasm density alters microtubule-bound *α*TAT1

We next aimed to understand the mechanisms by which cytoplasm density differentially regulates microtubule acetylation, because unlike the other tubulin PTMs analyzed, acetylation levels were disparately affected by increasing or decreasing crowdedness (Fig. 1). Acetylation of *α*-tubulin preferentially occurs on polymerized microtubules [40, 42, 43]. We therefore first examined if there was any difference in the soluble versus polymerized tubulin pools in response to osmotic challenges (Fig. S3). By fractionating soluble and polymerized *α*-tubulin, we determined the ratio of polymerized *α*-tubulin to total *α*-tubulin [55]. Cells exposed to different osmotic treatments showed a similar ratio of polymerized *α*-tubulin (*∼*45%), whereas nocodazole treatment decreased the ratio to *∼*5% and taxol treatment increased the ratio to *∼*99%, as expected (Fig. S3A). Consistent with this, we did not observe significant changes in overall microtubule intensity within cells exposed to different osmotic treatments (Fig. S3B-C). Next, we examined how changes in osmolarity and cytoplasm density affected proteins levels and the activity of the enzymes that control tubulin acetylation: *α*-tubulin acetyl transferase-1 (*α*TAT1: acetylating) and HDAC6 (deacetylating) [2]. HDAC6 protein levels remained unchanged in HeLa, BEAS-2B and RPE-1 cells in response to osmotic challenges, while *α*TAT1 levels were unchanged in HeLa cells and undetectable in BEAS-2B and RPE-1 cells via immunoblotting under our conditions (Fig. 4A-D).

**FIG. 4.**
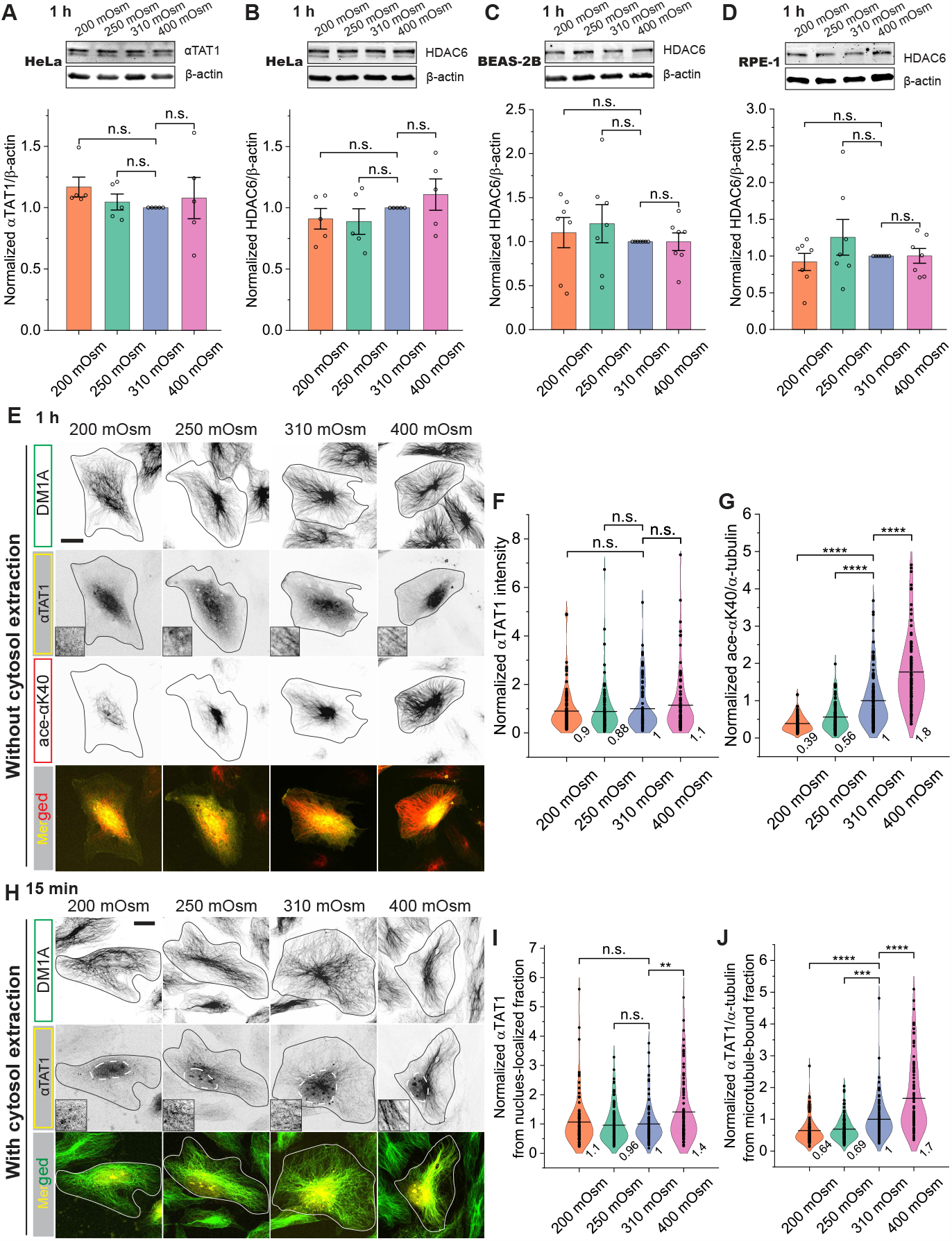
Cytoplasm crowdedness tunes microtubule-bound *α*TAT1. (A-D) WBs and corresponding quantification showing the relative changes of *α*TAT1 and HDAC6 in response to osmotic treatments for 1 hr in HeLa, BEAS-2B and RPE-1 cells compared to the loading control, *β*-actin. No significant changes were observed. HeLa *α*TAT1: n = 5; HeLa HDAC6: n = 5; BEAS-2B HDAC6: n = 7; RPE-1 HDAC6: n = 7. (E-G) Representative images and quantification comparing *α*TAT1 intensity and Ace-*α*K40/*α*-tubulin for cells transiently expressing *α*TAT1-mScarlet at reduced levels in response to different osmotic challenges for 1 hr. Cells were fixed with cold methanol immediately after the treatment and the cell cytosol was not extracted. Insets show a magnified portion of the images in which *α*TAT1-mScarlet is diffusive in cells treated with hypotonic solutions (200 and 250 mOsm). Microtubule-bound *α*TAT1 is difficult to quantify in cells treated with iso- and hypertonic solutions (310 and 400 mOsm) due to the high-background signal in the cytosol. n = 110, 95, 105 and 96 cells from 2 independent experiments. *α*TAT1 intensity: no significant difference; *α*K40/*α*-tubulin: \*\*\*\**p <* 0.0001. (H-J) Representative images and quantification comparing nuclear-localized and microtubule-bound fractions of *α*TAT1 for cells transiently expressing *α*TAT1-mScarlet at reduced levels in response to short-time (15 min) osmotic treatments. To measure the nuclear-localized and microtubule-bound fractions of *α*TAT1, the cytosol was extracted (PEM buffer with 1% TritonX-100 and 10 *μ*M taxol) before fixation with cold methanol. White dashed lines indicate the position of nuclear. Insets show a magnified portion of the images in which microtubule-bound *α*TAT1 is clearly visible after cytosol extraction especially for cells with hypertonic treatment. n = 93, 93, 88 and 87 cells from 2 independent experiments. Nuclear-localized fraction: \*\**p* = 0.0033; microtubule-bound fraction: \*\*\*\**p <* 0.0001, \*\*\**p* = 0.0001. Scale bars: 20 *μ*m. All graphs display all data points with means. Error bars indicate s.e.m. All *p*-values were calculated using a student’s t-test.

Osmotic shifts result in changes in cell volume [30, 35], and hence the intracellular concentration of enzymes (and substrates) if protein expression levels remain unchanged. We observed a minor decrease in the cell projection area upon hypertonic treatment (400 mOsm) and a small increase in the cell projection area upon hypotonic treatment (200 mOsm) (Fig. S3D). We did not detect a difference in cell projection area for the 250 mOsm condition, although this condition also caused a dramatic decrease in acetylation similar to the 200 mOsm treatment (Figs. S3D and 1C-D). We were therefore unconvinced that cell volume alone could account for the large differences in acetylation levels that we observed. To further test this idea, we transiently expressed mScarlet-tagged full length *α*TAT1 in BEAS-2B cells and subjected the cells to different osmotic challenges (Fig. 4E). In mammalian cells, the subcellular localization of *α*TAT1 is elusive, depending on the isoform and cell type examined, though it is rarely found to be associated with microtubules [40, 54, 56–58]. In BEAS-2B cells, *α*TAT1-mScarlet was mostly diffusive, but showed a weak-microtubule-binding pattern only in control and hyperosmotic conditions (Fig. 4E). While the fluorescence intensity of *α*TAT1 showed no significant changes in response to osmotic treatments (Fig. 4F), acetylation levels drastically changed, similar to the pattern observed without exogenous *α*TAT1 expression (Figs. 4G and 1F), suggesting that dilution of *α*TAT1 in the hypoosmotic condition is not likely the cause of the osmotic regulation of acetylation.

We next wanted to specifically analyze microtubule-bound *α*TAT1-mScarlet following osmotic treatment. We first subjected *α*TAT1-mScarlet-expressing cells to a brief, 15 min osmotic treatment, a time point at which acetylation is slightly increased in the hyperosmotic condition and unaffected in hypoosmotic conditions (Fig. 1I). Then, we extracted the cytosol as previously described [59], and fixed and stained the cells for microtubules. Remarkably, *α*TAT1-mScarlet signal was more evident along microtubules in the isoosmotic and hyperosmotic conditions when compared to cells without cytosol extraction. A considerable amount of *α*TAT1-mScarlet was found localized within the nucleus, similar to the localization of an alternative *α*TAT1 isoform in HeLa cells (Fig. 4H) [56]. We quantified the intensity of these two fractions and found that the nuclear-localized *α*TAT1-mScarlet increased 1.4-fold upon hypertonic treatment but remained unchanged upon hypotonic treatment (Fig. 4H-I). However, microtubule-associated *α*TAT1-mScarlet increased 1.7-fold upon hypertonic treatment and decreased 31% (250 mOsm) and 36% (200 mOsm) upon hypotonic treatments. The changes in microtubule-associated *α*TAT1-mScarlet following the osmotic shifts strongly suggests that the regulation of acetylation by cytoplasm density might be achieved by modulating the binding of *α*TAT1 to microtubules, which is inhibited in hypoosmotic conditions and promoted in hyperosmotic conditions.

### Microtubule lattice spacing regulates acetylation

To probe at the molecular mechanisms by which cytoplasm density dictates the *α*TAT1-microtubule interaction, we purified recombinant full length *α*TAT1-mScarlet (Fig. 5A) to analyze its binding and enzymatic activity in *ex vivo* assays. The observed size of *α*TAT1-mScarlet was 70 kDa by both SDS-PAGE and mass photometry analysis, with *∼*20% of the *α*TAT1-mScarlet population existing as a dimer (Fig. 5A-B). We tested its *ex vivo* acetylation efficiency on the microtubule cytoskeleton in detergent extracted BEAS-2B cells [59]. In the presence of acetyl coenzyme A (Acetyl-CoA), incubation with purified *α*TAT1-mScarlet caused an 8.4-fold increase in acetylation signal compared to untreated control cells (Fig. 5C-D). Next, we subjected cells to a 15 min osmotic challenge followed by the *ex vivo* acetylation assay. Before *ex vivo* acetylation, 15 min osmotic challenges did not significantly change normalized acetylation levels (Fig. 5E-F). Following acetylation by recombinant *α*TAT1-mScarlet, the normalized acetylation levels decreased by 27% (200 mOsm) and 12% (250 mOsm) after hypotonic pre-treatments (Fig. 5E-F), indicating that exogenous *α*TAT1-mScarlet activity was attenuated after hypoosmotic conditions. In contrast, when we exogenously expressed EGFP-MAP7 in cells prior to extraction, exogenous *α*TAT1-mScarlet strongly acetylated microtubules after all osmotic treatments (Fig. 5G-H), suggesting that MAP7 binding to microtubules promoted *α*TAT1 microtubule binding, enzymatic activity, or both. The interdimer distance within the microtubule lattice can be expanded or compacted depending on several factors. First, the nucleotide state of *β*-tubulin is thought to modulate interdimer packing. The GTP-bound state is correlated with an expanded interdimer distance, but hydrolysis of GTP to GDP is thought to cause compaction of the interdimer distance by *∼*2 angstroms [60, 61]. However, structural studies on non-mammalian microtubule lattices have revealed that yeast and worm microtubules exist in an expanded state regardless of nucleotide state [62, 63]. Second, MAP association alters the interdimer packing regardless of the nucleotide state of the tubulin dimers, with tau and MAP2 inducing and maintaining a compacted lattice [11], and motors such as kinesin-1 inducing lattice expansion [64, 65]. Third, drugs such as taxanes stabilize the expanded conformation of the lattice [11, 60]. In order to determine if different osmotic challenges may have altered the packing state of the microtubule lattice in our experiments, we analyzed the binding of the taxane, SiR-tubulin, which labels the expanded lattice [11, 66]. We found that in BEAS-2B cells, SiR-tubulin intensity increased 2.4-fold after hyperosmotic treatment and decreased 53% after hypoosmotic treatment (Fig. 5I-J). Taken together, these results indicate that increasing cellular density promotes microtubule lattice expansion, MAP7 association, and *α*TAT1 acetylation activity, whereas decreasing cellular crowdedness promotes lattice compaction, disfavoring MAP7 association and *α*TAT1-driven acetylation. Therefore, we conclude that the underlying microtubule lattice structure changes in response to osmotic challenges, providing a molecular rationale for the differential regulation of microtubule acetylation observed in response to changes in cytoplasmic density.

**FIG. 5.**
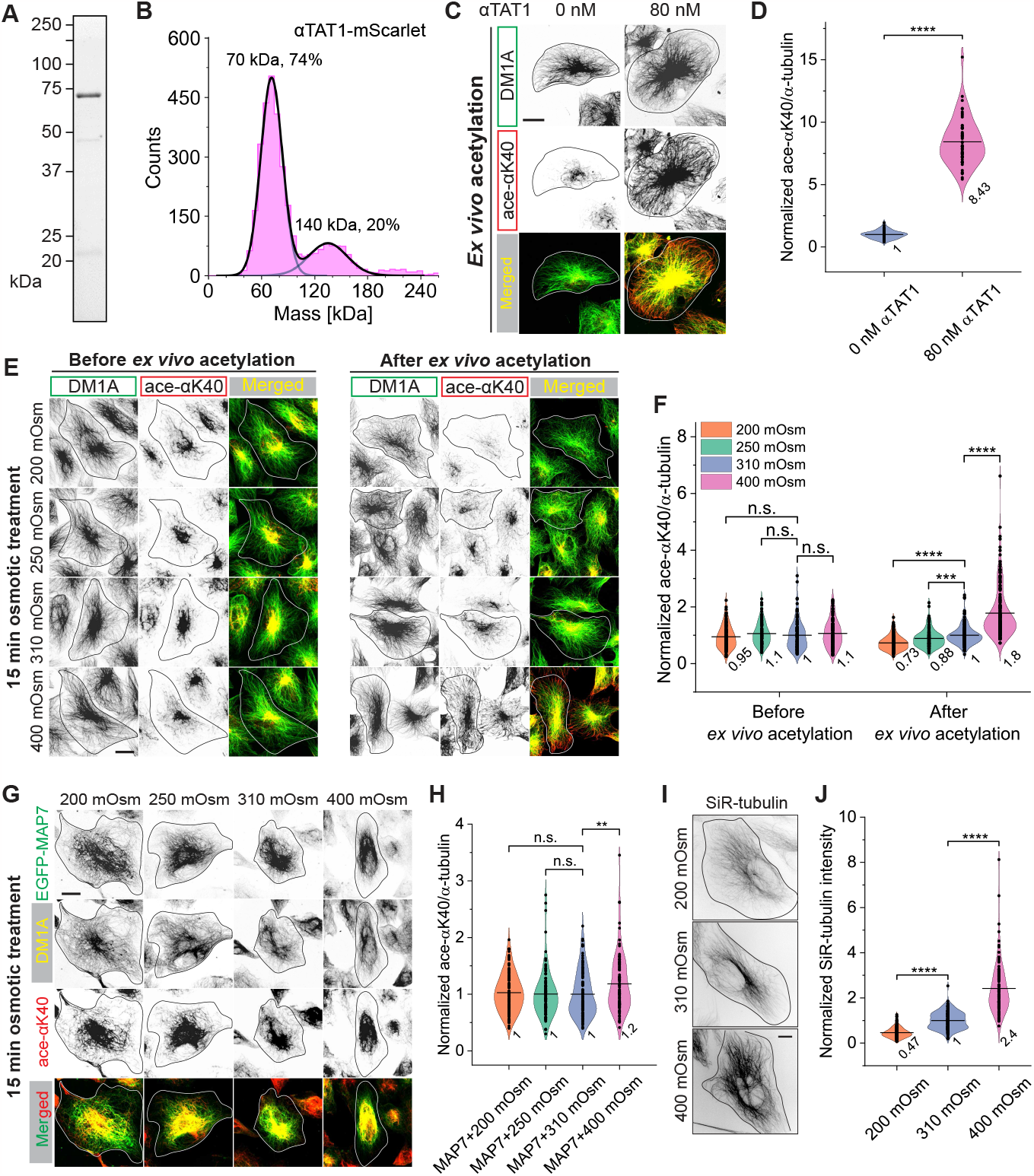
Microtubule lattice spacing regulates acetylation. (A) Stain-free precast SDS-PAGE gel of *α*TAT1-mScarlet purified from BL21-RIPL cells. (B) Mass photometry of *α*TAT1-mScarlet (expected mass: 65 kDa). Fits of mean mass values for monomer (observed mass: 70 kDa) and dimer (observed mass: 140 kDa) species, and relative fraction of particles with indicated mass, are shown. (C) Images of BEAS-2B cells following *ex vivo* acetylation. Cells were cytosol-extracted in the presence of taxol, incubated for 8 min without (left panels, 0 nM) or with 80 nM recombinant *α*TAT1 (right panels), fixed and stained for total tubulin (green) and acetylated *α*-tubulin (red). (D) Quantification of the relative changes of Ace-*α*K40 to total *α*-tubulin following *ex vivo* acetylation. n = 56 and 54 cells from 2 independent experiments, \*\*\*\**p <* 0.0001. (E) Images of BEAS-2B cells that were first treated with different osmotic solutions for 15 min and then either subjected (right panels) or not subjected (left panels) to *ex vivo* acetylation in the presence of 20 nM *α*TAT1. (F) Quantification of the *ex vivo* acetylation efficiency of microtubules in response to different osmotic pre-treatments for 15 min. Before *ex vivo* acetylation: n = 106, 93, 106 and 101 cells from 2 independent experiments; after *ex vivo* acetylation: n = 157, 165, 190 and 174 cells from 4 independent experiments, \*\*\*\**p <* 0.0001, \*\*\**p* = 0.00076. (G) Images of BEAS-2B cells transiently expressing EGFP-MAP7 that were first treated with different osmotic solutions for 15 min and then subjected to *ex vivo* acetylation in the presence of 20 nM *α*TAT1. (H) Quantification of the *ex vivo* acetylation efficiency of microtubules in response to different osmotic pre-treatments for 15 min upon MAP7 overexpression in cells. n = 122, 100, 114 and 92 cells from 3 independent experiments, \*\**p* = 0.0038. Scale bars: 20 *μ*m. (I) Images of BEAS-2B cells labeled with SiR-tubulin (0.5 *μ*M) showing the distribution of SiR-tubulin following osmotic treatments for 15 min. Scale bar: 10 *μ*m. (J) Quantification of the relative changes of SiR-tubulin intensity in response to different osmotic treatments. n = 163, 221 and 108 cells from 3 independent experiments, \*\*\*\**p <* 0.0001. All graphs display all data points with means. All *p*-values were calculated using a student’s t-test.

### MAP7 excludes VASH1 but not *α*TAT1 *in vitro*

Hypoosmotic conditions induced microtubule detyrosination and the disassociation of MAP7 from microtubules (Figs. 1-2). In contrast, hyperosmotic conditions increased MAP7 association with microtubules and promoted acetylation (Figs. 1-2). We were therefore curious how the presence of MAP7 on microtubules affects the enzymes responsible for these two modifications. Detyrosination of *α*-tubulin is catalyzed by an enzymatic complex composed of a vasohibin (VASH1 or VASH2) and a small vasohibin-binding protein (SVBP) [2]. Using TIRF microscopy with purified proteins, we examined the binding of VASH1-SVBP on microtubules in the absence and presence of MAP7, and found that microtubule binding of VASH1-SVBP was completely inhibited by MAP7 (Fig. 6A). In contrast, we found that *α*TAT1 microtubule-binding was only mildly reduced by the same concentration of MAP7 (Fig. 6B). We analyzed published cryo–electron microscopy (cryo-EM) structures of microtubules decorated with MAP7 [67] or VASH1-SVBP [68], and observed a steric clash between the MAP7 and VASH1 binding sites on *α*-tubulin (Fig. 6C). Therefore, the presence of MAP7 on microtubules may interfere with the detyrosination process by physically occluding VASH1 from the lattice (Fig. 6C). The observed increase in detyrosination in hypoosmotic conditions may be due to dissociation of MAP7, thereby facilitating VASH1-SVBP association with microtubules. *α*TAT1 binds inside the microtubule lumen and therefore does not have overlapping binding sites with MAP7 [44]. Interestingly, in analyzing the cryo-EM structures of tubulin from a MAP7-decorated lattice and tubulin from GDP-, taxol- or GMPCPP-lattices, we noticed that the *α*K40-loop (a.a. 38-46) containing the *α*TAT1 acetylation site, K40, was ordered in the MAP7-bound configuration but not in the GMPCPP-structure (Fig. 6D; GDP and taxol lattices do not have ordered *α*K40-loops either). These results suggest that MAP7 binding may facilitate the acetylation process through allosterically-induced ordering of the luminal *α*K40-loop.

**FIG. 6.**
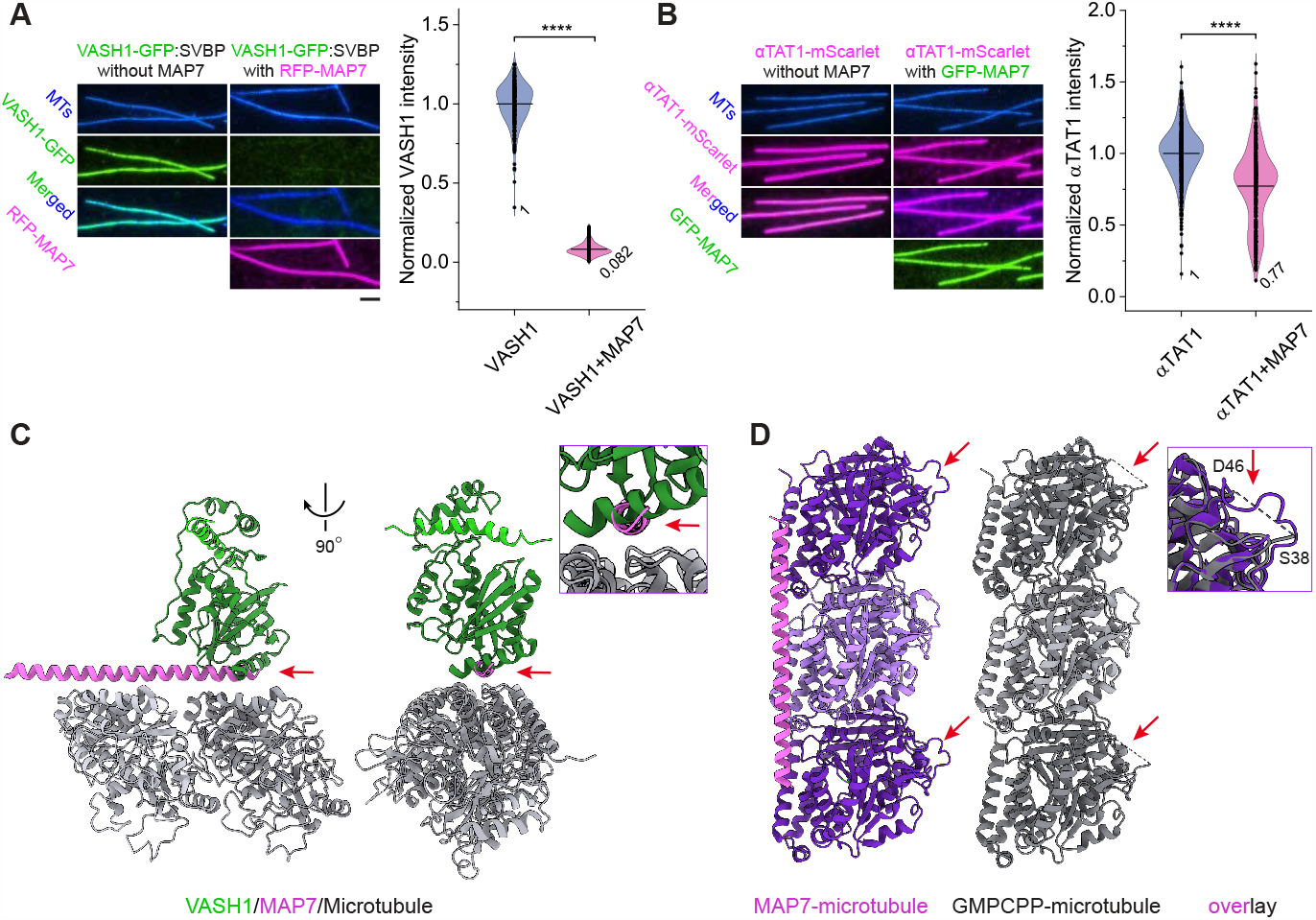
MAP7 excludes VASH1 but not *α*TAT1 *in vitro*. (A) Representative fluorescent images and quantification showing VASH1-GFP (100 nM) signals along microtubules in the absence of RFP-MAP7 and presence of 50 nM RFP-MAP7. VASH1-GFP and SVBP were co-purified from insect cells using a dual expression vector. Scale bar: 2 *μ*m. n = 239 and 291 microtubules from 2 independent experiments, \*\*\*\**p <* 0.0001. (B) Representative fluorescent images and quantification showing *α*TAT1-mScarlet (25 nM) signals along microtubules in the absence of GFP-MAP7 and presence of 25 nM GFP-MAP7. Scale bar: 2 *μ*m. n = 434 and 432 microtubules from 3 independent experiments, \*\*\*\**p <* 0.0001. (C) Overlay of cryo-EM structures of microtubules (gray) decorated with MAP7 (magenta) or VASH1-SVBP (green). Inset shows a magnified image of the clash between the MAP7 binding site and the VASH1 contact site on *α*-tubulin. Red arrows indicate the clash sites. (D) Cryo-EM structures of MAP7-decorated (purple) and GMPCPP-(gray) microtubules lattice and their overlay. Red arrows indicate the *α*K40-loops. Inset shows a magnified image of the *α*K40-loops. All graphs display all data points with means. All *p*-values were calculated using a student’s t-test.

### Cytoplasm crowdedness differentially regulates cargo transport

We have demonstrated that cells respond to different cytosolic densities by altering tubulin PTMs, MAP association, and the underlying microtubule lattice state. But why? Prior studies have described roles for tubulin PTMs, MAPs, and lattice spacing in directing motor transport [21, 53, 69–73]. We therefore investigated the impact of these altered microtubule landscapes on cargo transport. We examined the transport of two specific organelle types in response to alterations in cytoplasmic density: secretory or dense core vesicles (EGFP-Rab6A) that are primarily transported anterogradely from the perinuclear region to cell periphery by kinesin-1 and kinesin-3 motors [73], and early endosomes (EGFP-Rab5) that are predominantly transported in the retrograde direction by dynein-dynactin complexes (Fig. S4) [74, 75]. The movement of vesicles can be divided into two distinct factions: the immobile tracks with no obvious directed motion during the 3 min observation time window, and the mobile tracks with persistent directed motion (Figs. 7A and S4). Individual motile trajectories displayed switching between states of diffusive movement or “pauses” and states of directed “runs” that were attributed to active motor-driven transport along microtubules (Fig. 7B), consistent with prior cellular studies on vesicle transport [73, 75–77]. We used a previously developed algorithm to extract the distinct motion states and analyzed the mobile ratio of vesicles, the ratio of run time to total travel time for mobile vesicles, the velocities of individual run segments, and their run times [76]. The obtained MSDs of each run segment revealed a parabolic function indicative of active transport (Fig. 7C). These 4 parameters thus gave us an in-depth, unbiased characterization of the full spectrum of the motions observed.

**FIG. 7.**
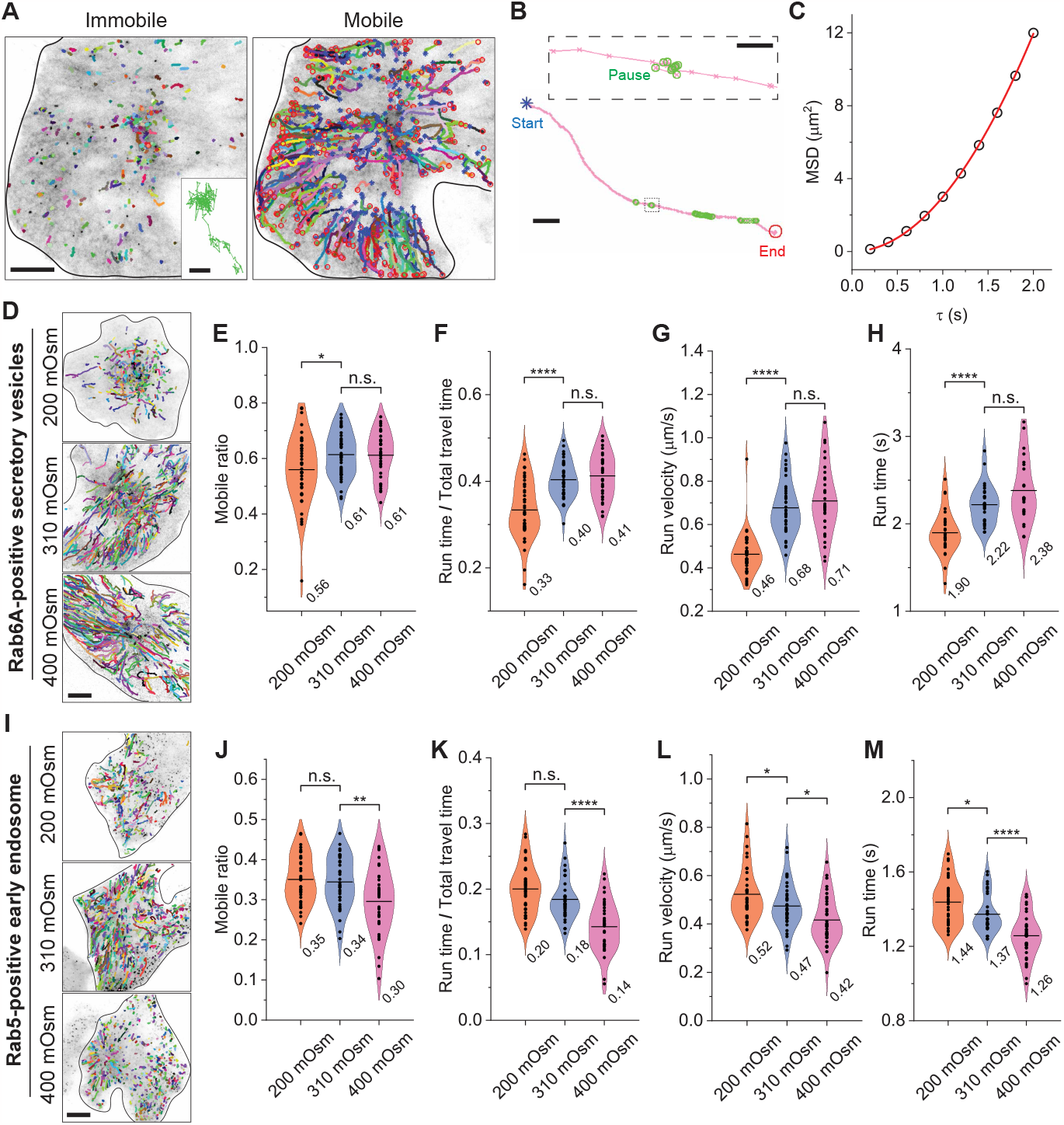
Cytoplasm crowdedness differentially tunes cargo transport. (A) Representative trajectories (colored lines) of EGFP-Rab6A-positive secretory vesicles in a BEAS-2B cell (10 fps for 3 min) showing the relative immobile (left) and mobile (right) fractions of all movements observed. Each color represents an individual trajectory. For the mobile fraction, blue stars and red circles mark the start and end point of each trajectory, respectively. Rab6A-positive vesicles are mostly transported towards the plus end of microtubules. Scale bar: 10 *μ*m. Inset shows a magnified view of an immobile vesicle trajectory. Scale bar: 100 nm. (B) A magnified trajectory image showing a Rab6A-positive vesicle undergoing a mixed mode of run and pause motions. Pause states are highlighted by green circles. Motion states were detected and extracted by a homemade algorithm [76]. Scale bar: 2 *μ*m. Inset shows a magnified view of the pause state. Scale bar: 200 nm. (C) Measured MSD as a function of delay time *τ* for the run segment of a representative trajectory. Run velocity *v* of each run segment is obtained by fitting the data to the parabolic function (red solid line) with *MSD*(*τ*) = *v*^2^*τ* ^2^. (D) Representative trajectories of the mobile fraction of Rab6A-positive vesicles for cells treated with different osmotic solutions. EGFP-Rab6A expressing cells were imaged after incubation with corresponding solutions. Scale bar: 10 *μ*m. (E-H) Measured mobile ratios, run time/total travel time ratios, run velocities, and run times of Rab6A-positive vesicles showing the changes of cargo transport in response to different osmotic treatments. Mobile ratio, \**p* = 0.023; run time/total travel time, \*\*\*\**p <* 0.0001; run velocity, \*\*\*\**p <* 0.0001; run time, \*\*\*\**p <* 0.0001. The total trajectories analyzed from the mobile fraction are n = 29688, 32248 and 22572 from n = 42, 40 and 34 cells from 3 independent experiments. Each data point is the average of the measured quantities from individual trajectories from a single cell. (I) Representative trajectories of the mobile fraction of Rab5-positive endosomes for cells treated with different osmotic solutions. EGFP-Rab5 expressing cells were imaged after incubation with corresponding solutions. Scale bar: 10 *μ*m. (J-M) Measured mobile ratios, run time/total travel time ratios, run velocities, and run times of Rab5-positive endosomes showing the changes of cargo transport in response to different osmotic treatments. Mobile ratio, \*\**p* = 0.0070; run time/total travel time, \*\*\*\**p <* 0.0001; run velocity, \**p* = 0.048 and 0.015; run time, \**p* = 0.017, \*\*\*\**p <* 0.0001. The total trajectories analyzed from the mobile fraction are n = 33130, 28305 and 25248 from n = 37, 34 and 36 cells from 3 independent experiments. Each data point is the average of the measured quantities from individual trajectories from a single cell. All graphs display all data points with means. All *p*-values were calculated using a student’s t-test.

Interestingly, under the hypoosmotic condition (200 mOsm), the Rab6A-vesicle mobile ratio, run time/total travel time ratio, run velocity and run time were all decreased, whereas these parameters were unchanged for the hyperosmotic condition (400 mOsm) when compared to the isotonic control (Fig. 7D-H). The same analysis performed on Rab5-positive early endosomes found that all four parameters decreased in the hyperosmotic condition, while the run velocity and run time showed significant increases in the hypoosmotic condition compared to the isotonic control (Fig. 7I-M). Taken together, these results indicate that decreasing cellular crowdedness selects against anterograde transport of secretory vesicles in favor of retrograde transport of early endosomes, while increasing cellular crowdedness selects against retrograde transport of early endosomes. We hypothesize that the distinct transport behaviors observed for Rab6-vesicles and Rab5-positive early endosomes in response to different osmotic treatments is not simply due to changes in the physical properties of the cytoplasm, but may be attributed to the changes in MAP association and tubulin PTMs.

## Discussion

Our study presents an analysis of the mechanisms utilized by cells to tailor their microtubule cytoskeletons in response to changes in macromolecular crowding conditions. Increasing cytoplasmic density induces MAP7-bound, acetylated microtubules, while decreasing cytoplasmic density induces MAP7 release from microtubules and detyrosinated microtubules (Figs. 1-2). These distinct microtubule populations in turn differentially influence intracellular transport by molecular motors. We hypothesize that cells specifically utilize MAPs to control tubulin PTMs and microtubule-based transport to efficiently adapt to challenging environmental conditions.

What could be the physiological significance of our observations? For cells to maintain a constant size, they must balance endocytosis and exocytosis [78–80]. In response to hypoosmotic conditions and decreased crowding, the cell responds by downregulating MAP7-association and acetylation. The functional consequence of these events is a substantial reduction in anterograde transport of Rab6a secretory vesicles. This response may therefore be restricting secretory vesicle transport and fusion to prevent additional membrane insertion and cell swelling [78, 79]. In response to hyperosmotic conditions and increased crowding, we observe a decrease in the retrograde transport of Rab5 early endosomes, possibly indicating a response leading to restriction of endocytosis in order to prevent cell shrinking. We therefore speculate that cells modify their microtubule landscape to tune cargo transport and regulate endocytosis and exocytosis to protect themselves against drastic changes in cell volume and membrane tension [81, 82].

To respond to changes in osmolarity, the cell may specifically target the microtubule binding properties of MAP7 for multiple reasons. First, MAP7 differentially affects tubulin PTMs. We have found that MAP7 promotes the acetylation of microtubules. Acetylated microtubules have been shown to be more flexible and more resistant to breakage [13], which may be advantageous to withstand the increased viscosity and rigidity of the hyperosmotic cytoplasm [35, 83]. Under hypoosmotic conditions, MAP7 dissociation correlates with an increase in microtubule detyrosination, perhaps due to an indirect augmentation of VASH-SVBP binding to the lattice. In this case, the cell may require a microtubule population marked by detyrosination to support the increased dynamicity that has been observed in hypoosmotic conditions [30]. Second, since MAP7-family proteins are required for the *in vivo* functions of kinesin-1 [8, 72, 84], their presence or absence will dictate kinesin-1 driven transport. Not only does MAP7 directly recruit kinesin1 to microtubules [8, 46, 85], but it protects acetylated microtubules, which are preferred by kinesin-1 [69, 86]. Therefore, MAP7 facilitates kinesin-1 motility both directly and indirectly. Increased viscosity or cytoplasmic density assumes a general role in slowing down motor protein transport both *in vitro* and *in vivo* [31–33]. Indeed, we find that an increase in macromolecular crowding significantly decreases retrograde transport. Surprisingly, it does not disrupt anterograde transport of Rab6a secretory vesicles (Fig. 7), leading us to conclude that the cell may enhance MAP7 binding to ensure kinesin1 cargo transport of these vesicles to plasma membrane persists in the presence of increased viscosity and drag forces. The opposite is true under lower density conditions: MAP7 dissociation results in a significant decline in the anterograde transport of secretory vesicles, which may be necessary for the cell to manage cell swelling as described above. Therefore, tuning MAP7 association with microtubules equips the cell with various strategies for adaptation to changes in macromolecular crowding and cell size.

We were surprised to observe a distinct decoupling of microtubule acetylation and detyrosination. Although both of these modifications are associated with “stable” microtubules [2, 13, 52] and often occur on the same subset of microtubules [52, 53, 86, 87], we have discovered that specific cellular conditions select for one PTM over the other, indicating that these modifications can be separated and utilized for specific cellular purposes. Our results are consistent with prior studies that show acetylation and detyrosination modifications are not always tightly linked, such as in cancer cells, in migrating cells, and upon microtubule regrowth after depolymerization [52, 88, 89]. Therefore, although these modifications are not mutually exclusive, we show here that MAP7 association has the ability to promote acetylation and limit detyrosination, thus decoupling them.

How does the cell control MAP7 association with microtubules? We have found that the underlying lattice state is altered in response to crowding conditions with a lower cytoplasmic density favoring compacted lattices and higher cytoplasmic density favoring expanded lattices. The expanded lattice correlates with increased binding of MAP7 and *α*TAT1, and acetylation. The compacted microtubules in the hypoosmotic condition disfavor MAP7 and *α*TAT1 binding, indicating these proteins preferentially bind expanded lattices. Considering kinesin-1 predominantly binds the expanded lattice [64, 90], it is logical that MAP7, the primary recruiter of kinesin-1, and *α*TAT1, the modifying enzyme that produces the preferred lattice of kinesin-1, also favor the expanded lattice. We hypothesize that MAP7 binding may drive lattice expansion because overexpression of MAP7 prior to hypotonic treatment appears to “lock” microtubules in a particular state that promotes *α*TAT1 association and activity. We previously showed that tau is a mechanosensitive MAP that actively compacts the expanded microtubule lattice [11]. MAP7 not only competes with tau, but displaces tau from the microtubule [8], consistent with MAP7 possibly facilitating lattice expansion. It will be interesting to explore how different cells employ these mechanosensitive MAPs in future studies.

Together, this work characterizes the interplay between the MAP code and the tubulin code in modulating the biophysical parameters of microtubules to allow them to withstand changes in cytoplasmic forces, and in directing the intracellular transport of specific cargoes to respond to osmotic pressure and macromolecular crowding. Overall, our data indicate that the cell utilizes its MAP toolbox to destine microtubules for precise functions in response to changes in its physical environment.

## Methods

Details about the materials and methods are given in Supplementary Materials.

## Data availability

All data generated or analyzed during this study are included in the manuscript or as source files.

## Acknowledgments

The authors wish to thank all the members of the Ori-McKenney and McKenney laboratories for their kind help and feedback. We thank Dr. Shinsuke Niwa (Tohoku University, Japan) for generously proving the recombinant VASH1-SVBP complex and Dr. Scott Hansen (University of Oregon) for providing the DeltaCMV vector for expressing proteins at reduced levels in mammalian cells. This work is supported by the NIH grant 1R35GM133688 to K.M.O.-M.

## Author Contributions

Y. S. and K. M. O. M. conceived the project and designed the experiments. K. M. O. M. performed the *in vitro* TIRF-M experiments on VASH1-SVBP and the structural analysis of MAP7 and VASH1-SVBP decorated microtubules. Y. S. performed all other experiments and analyzed the data. Y. S. and K. M. O. M. wrote the manuscript.

## Competing financial interests

The authors declare no competing financial interests.

## Notes

### Competing Interest Statement

The authors have declared no competing interest.

