## Supplementary Information for "Macromolecular Crowding Tailors the Microtubule Cytoskeleton Through Tubulin Modifications and Microtubule-Associated Proteins"

### Supplemental Materials concerning the article: Macromolecular Crowding Tailors the Microtubule Cytoskeleton Through Tubulin Modifications and Microtubule-Associated Proteins

Yusheng Shen, Kassandra M. Ori-McKenney
*Department of Molecular and Cellular Biology, University of California, Davis, Davis, CA 95616, USA*
(Dated: June 12, 2023)

#### I. MATERIALS AND METHODS

**Antibodies.** For Western blotting and immunostaining, we used rabbit monoclonal recombinant antibodies against $\alpha$-tubulin with acetylation at $\alpha$K40 (EPR16772, ab179484; Abcam), HDAC6 (D21B10; Cell Signaling), and DCLK1 (D2U3L; Cell Signaling), mouse monoclonal antibodies against $\alpha$-tubulin with acetylation at $\alpha$K40 (6-11B-1; Invitrogen), polyglutamylated tubulin (clone B3, T9822; Sigma), $\alpha$-tubulin (DM1A, T9026; Sigma), $\beta$-actin (AC74; Sigma), a chicken polyclonal antibody against GFP (A10262; Invitrogen), a mouse polyclonal antibody against MAP7 (H00009053-B01P; Abnova), rabbit polyclonal antibodies against detyrosinated $\alpha$-tubulin (AB3201; Sigma), MAP4 (PA5-35920; Invitrogen), $\alpha$TAT1 (HPA046816; Sigma), DCLK1 (ab31704; Abcam) and MAP9 (PA5-58145; Invitrogen), a rat monoclonal antibody against tyrosinated $\alpha$-tubulin (YL1/2, MA1-80017; Invitrogen) and a rat monoclonal recombinant antibody against GFP (3H9, ab252881; Abcam). The following secondary antibodies were used: DyLight$^{TM}$ 680 goat anti-mouse (35518, Invitrogen) and DyLight$^{TM}$ 800 goat anti–rabbit (SA5-10036, Invitrogen) for Western blotting and Alexa Fluor 488–, 555–, and 647–conjugated goat antibodies against rabbit, rat, mouse and chicken IgG (Invitrogen) for immunofluorescence.

**Cell culture, transfection and drug treatments.** BEAS-2B cells (ATCC, CRL-9609) and HeLa cells (ATCC, CRM-CCL-2) were maintained in Dulbecco's modified Eagle's medium (DMEM, Life Technologies), and RPE-1 cells (ATCC, CRL-4000) were maintained in Dulbecco's modified Eagle's medium nutrient mixture F-12 (DMEM/F12, Life Technologies). Both media were supplemented with 10% fetal bovine serum (FBS), 50 units/mL of penicillin and 50 $\mu$g/mL of streptomycin. All cell cultures were maintained in a 95% air/5% $CO_2$ atmosphere at 37 °C. The cell lines were routinely confirmed to test negative for mycoplasma contamination. For live cell imaging, cells were seeded at a density of approximately $1\times 10^4$ cm$^{-2}$on a glass coverslip which was placed in a 35-mm polystyrene tissue-culture dish.

The expression vectors used in this study were pCDNA3.1-pCMV-PfV-GS-Sapphire (Addgene plasmid 116933; for mammalian expression of 40 nm-GEMs), pDeltaCMV-sfGFP-MAP7 (full-length human MAP7 from GE Dharmacon MGC Collection BC025777), pDeltaCMV-sfGFP-MAP4 (full-length human MAP4 isoform 1 from the McKenney lab) [1], pEGFP-MAP7, pEGFP-MAP7-Cter (a.a. 354-749), EB1-tdTomato (Addgene plasmid 50825), pDeltaCMV-$\alpha$TAT1-mScarlet (human isoform 4 from Addgene plasmid 27099), pET28a-$\alpha$TAT1-mScarlet-StrepII (human isoform 4 from Addgene plasmid 27099), pEGFP-Rab6A (Addgene plasmid 49469) and pEGFP-Rab5 (Addgene plasmid 49888). MAP7, MAP4 and $\alpha$TAT1 proteins were cloned in frame using Gibson cloning into a pDeltaCMV vector (a gift from Dr. Scott Hansen for expressing proteins at reduced levels) with an N-terminal superfolder GFP (sfGFP) cassette or a C-terminal mScarlet cassette. MAP7 and its C-terminal truncation, MAP7-Cter, proteins were also cloned in frame using Gibson cloning into a pEGFP-C1 vector. $\alpha$TAT1 protein was also cloned in frame using Gibson cloning into a pET28a vector with a C-terminal mScarlet-strepII cassette. Transfections were performed by using the Lipofectamine 2000 reagent kit (Invitrogen) according to manufacturer's instructions. Cells were generally transfected for 8 h with 1 $\mu$g plasmids when the density reached ∼80% confluency.

To inhibit HDAC6 activity, cells were treated with tubacin (5 $\mu$M, Sigma) in culture medium for 1 h before being fixed. The control group was treated with DMSO. To either disrupt or stabilize the microtubules, cells were treated with nocodazole (1.5 $\mu$M, Sigma) or taxol (5 $\mu$M, Sigma) in culture medium for 1 h, respectively, before being lysed to fractionate the soluble and polymerized $\alpha$-tubulin.

**Osmotic treatments.** To manipulate the cytoplasm crowdedness, cells were treated with and imaged in extracellular osmotic environments ranging from hypoosmotic to hyperosmotic conditions (200–400 mOsm). These solutions were prepared by adding, respectively, 0.28 (∼400 mOsm, hypertonic), 0.2 (∼310 mOsm, isotonic control), 0.15 (∼250 mOsm, hypotonic) and 0.1 M (∼200 mOsm, hypotonic) mannitol to a hypotonic base solution (in millimolar: 40 NaCl, 5 KCl, 1 $CaCl_2$, 2 $MgCl_2$, 10 HEPES (pH 7.4), ∼91 mOsm) to maintain a constant ionic strength.

**Western blotting.** Total BEAS-2B, HeLa and RPE-1 cell lysates were prepared in RIPA buffer containing 50 mM Tris-HCl, pH 7.4, 150 mM NaCl, 1% Triton X-100, 0.5% sodium deoxycholate, 0.1% SDS, protease inhibitor cocktail (Roche) and 1 mM PMSF. All immunoblots were developed with corresponding primary antibodies

(1:2000) and fluorescent secondary antibodies (1:4000), and visualized by infrared laser scanner (Odyssey CLx, Licor). Protein bands in Western blots were quantified using Fiji software (https://fiji.sc/).

**Fractionating soluble and polymerized -tubulin.** We followed the published protocols to fractionate soluble and polymerized tubulin [2]. Briefly, cells were first incubated with soluble tubulin extraction buffer A (137 mM NaCl, 20 mM Tris-HCl, pH 7.4, 1% Triton X-100, and 10% glycerol) at 4 °C for 1 min, and the buffer was then collected and saved as the soluble fraction. Next, polymerized tubulin extraction buffer B (A + 1% SDS) was immediately added to cells. After incubation for 1 min, cells were scraped, and the polymerized fraction was collected, sonicated briefly and incubated on ice for 30 min. The soluble and polymerized fraction were then used for Western blotting. To prove the efficiency of the extraction of soluble and polymerized tubulin, cells treated with nocodazole or taxol were used as controls.

**Immunostaining and confocal microscopy.** For immunofluorescence cell staining, BEAS-2B cells were fixed in −20 °C methanol for 10 min, and then blocked with 4% bovine serum albumin (Sigma-Aldrich) in PBS at room temperature for 2 h. Next, cells were incubated with primary antibodies against acetylated tubulin (1:200), detyrosinated tubulin (1:200), tyrosinated tubulin (1:500), polyglutamylated tubulin (1:200), $\alpha$-tubulin (1:500), MAP7 (1:100), MAP4 (1:200), DCLK1 (1:100), MAP9 (1:200) and GFP (1:200), and corresponding secondary antibodies (1:200), each for 1 h. Cells were washed with PBS (5 min each for three times) before and after the incubation with secondary antibodies, and then mounted onto a microscope slide with ProLong$^{TM}$ Gold Antifade Mountant (Invitrogen), and then examined by using spinning disk confocal microscopy. The spinning disk confocal was performed on an inverted research microscope Eclipse Ti2-E with the Perfect Focus System (Nikon), equipped with a Plan Apo 60× NA 1.40 oil objective, a Crest X-Light V3 spinning disk confocal head (Crest-Optics), a Celesta light engine (Lumencor) as the light source, a Prime 95B 25MM sCMOS camera (Teledyne Photometrics) and controlled by NIS elements AR software (Nikon). For live cell imaging of SiR-tubulin labelled microtubules, a live cell imaging chamber (H301-Nikon-TI-S-ER, Oko Labs) was equipped to the microscope to provide optimal culture conditions (95% air/5% $CO_2$ atmosphere at 37 °C) for cells during imaging. After SiR-tubulin labelling, the cell-containing glass coverslip was mounted on a coverslip holder (SC15012, Aireka Cells), which was finally mounted on the microscope. The fluorescent images for BEAS-2B cells were collected over a stack of vertical z-sections across the entire cell ∼4 $\mu$m thickness. The final fluorescent images and their fluorescent intensities shown in the main text are based on the z-averaged images by using Fiji software (https://fiji.sc/).

**Protein expression and purification.** Full length human $\alpha$TAT1 (isoform 4) was expressed in BL21-RIPL bacterial cells with a C-terminal mScarlet-strepII fusion. Cells were grown at 37 °C until an optical density at 600nm (O.D. 600) of ∼0.6 and protein expression was induced with 0.5 mM IPTG for 16 h at 16 °C before they were harvested and frozen [3]. Cells were thawed at 37 °C and resuspended in lysis buffer (50 mM Tris pH 8, 150 mM K-acetate, 2 mM Mg-acetate, 1 mM EGTA, 10% glycerol) with protease inhibitor cocktail (Roche), 1 mM DTT, 1 mM PMSF, and DNAseI on ice. Next, cells were passed through an Emulsiflex (Avestin) press and cleared by centrifugation at 23,000g for 20 mins. Proteins were first affinity purified from the clarified lysate using Streptactin XT beads (Qiagen/IBA) and further purified by cation exchange column 5 mL HiTrapS (GE Healthcare) in HB buffer (35 mM HEPES pH 7.5, 1 mM $MgCl_2$, 0.2 mM EGTA and 0.1 mM EDTA) and eluted with 0–1 M NaCl gradient over 20 coloum volumes. Fractions were analyzed by SDS polyacrylamide gel electrophoresis (SDS-PAGE), and then collected, combined, stored into single use aliquots and flash frozen in liquid nitrogen.

For *in vitro* experiments, RFP-MAP7 [4] and GFP-MAP7 [4] were prepared as previously described. Human VASH1-GFP:SVB complex is a kind gift from Dr. Shinsuke Niwa (Tohoku University, Japan).

**Microtubule assembly.** Tubulin was isolated from porcine brain using the high-molarity PIPES procedure as previously described [5], and then labelled with biotin NHS ester and Dylight-405 NHS ester as previously described [6]. Microtubules were assembled by incubating a mixture of native tubulin, biotin-tubulin, and fluorescent-tubulin (∼10:1:1 ratio, total ∼100 $\mu$M) in BRB80 buffer (80 mM PIPES, 1 mM $MgCl_2$, 1 mM EGTA, pH 6.8 with KOH) with 1 mM GTP for 15 min at 37 °C, and then stabilized with 20 $\mu$M taxol for 20 min. Microtubules were pelleted 20,000 g by centrifugation over a 25% sucrose cushion in BRB80 buffer to remove unpolymerized tubulin, and then resuspended and kept in 50 $\mu$L BRB80 buffer containing 10 $\mu$M taxol.

**Mass photometry.** To prepare imaging chambers for mass photometry, high precision glass coverslips (No. 1.5H, Cat 0107222, Marienfeld) were cleaned by sequential sonication in ultrapure $H_2O$, isopropanol and ultrapure $H_2O$, each for 10 min, and then dried with filter air. Culturewell$^{TM}$ gaskets (Cat 103250, Grace Bio-Labs) were then cut, sequentially rinsed with isopropanol and ultrapure $H_2O$, dried with filtered air, and placed onto the freshly cleaned coverslips. To focus the objective on the glass-buffer interface, 15 $\mu$L SRP90 buffer (90 mM HEPES-KOH pH 7.4, 50 mM K-acetate, 2 mM Mg-acetate, 1 mM EGTA, 10% glycerol) was added to the well, and the focal position was identified and secured in place with an autofocus system based on total internal reflection for the entire measurement. Immediately prior to mass photometry measurements, protein stocks were diluted directly in SRP90 buffer with typical working concentrations ∼5–20nM, and 5 $\mu$L of diluted proteins were introduced to the imaging well and imaged at the rate of 1 kHz for 60 s following autofocus stabilization. All data were acquired using an OneMP mass photometer (Refeyn Ltd) that was controlled by AcquireMP (Refeyn Ltd). Each sample was measured at least three times independently. Calibration standard was generated using bovine serum albumin (BSA, Sigma), Alcohol dehydrogenase (A7011, Sigma) and beta-amylase (A8781, Sigma). Data analysis was performed using DiscoverMP (Refeyn Ltd).

**Total internal reflection fluorescence microscopy.** TIRF microscopy experiments were performed on an inverted research microscope Eclipse Ti2-E with the Perfect Focus System (Nikon), equipped with a 1.49 NA 100× TIRF objective with the 1.5× tube lens setting, a Ti-S-E motorized stage, piezo Z-control (Physik Instrumente), LU-N4 laser units (Nikon) as the light source, an iXon DU897 cooled EMCCD camera (Andor) with an high-speed emission filter wheel (ET480/40M for mTurquoise2, ET525/50M for GFP, ET520/40M for YFP, and ET632/60M for mRuby2; Chroma). The microscope was controlled with NIS Elements software (Nikon). All live cell experiments were performed in a live cell imaging chamber (H301-Nikon-TI-S-ER, Oko Labs) that was equipped to the microscope to provide optimal culture conditions (95% air/5% $CO_2$ atmosphere at 37 °C) for cells during imaging. After cell transfection, the cell-containing glass coverslip was mounted on a coverslip holder (SC15012, Aireka Cells), which was finally mounted on the microscope. All *in vitro* experiments were conducted at room temperature. The TIRF flow chambers were assembled from acid-washed coverslips as described previously (http://labs.bio.unc.edu/Salmon protocolscoverslippreps.html), pre-cleaned slide and double-sided sticky tape. To immobilize microtubules in the chamber, chambers were first incubated with 0.5 mg/mL PLL-PEG-biotin (Surface Solutions) for 5 min, followed by 0.5 mg/mL streptavidin for 5 min. Microtubules were diluted into BC buffer (80 mM PIPES pH 6.8, 1 mM $MgCl_2$, 1 mM EGTA, 1 mg/mL BSA, 1 mg/mL casein) supplemented with 10 $\mu$M taxol, flowed into the chambers and incubated for 10 min to allow biotin-labelled microtubules to adhere to the streptavidin-coated surface. Unbound microtubules were washed away with TIRF assay buffer (90 mM HEPES-KOH pH7.4, 50 mM K-acetate, 2 mM Mg-acetate, 1 mM EGTA, 10% glycerol, 0.5% Pluronic F127, 0.1 mg/mL biotin-BSA, 0.2 mg/mL -casein and 10 $\mu$M taxol). All *in vitro* experiments with MAP7, VASH1-SVBP and $\alpha$TAT1 were performed in TIRF assay buffer, and were quantified by pooling data from at least two chambers performed on multiple days by using Fiji software (https://fiji.sc/).

**Single particle tracking and analysis.** Single-particle tracking (SPT) was performed using a home-made tracking program written in MATLAB as previously described [7], which is based on the standard tracking algorithm [8, 9]. With this advanced SPT algorithm, we were able to obtain the position $\mathbf{r}(t)$ at time $t$ for GEMs, Rab6A-positive secretory vesicles and Rab5-positive early endosomes, and their trajectories were constructed from the consecutive images. To study the diffusion dynamics of GEMs, we first selected the mobile trajectories from the whole set of GEMs trajectories. This is achieved by computing the radius of gyration $R_g(\tau)$ of each GEMs trajectory obtained over a time period of $\tau$,

$$R_g^2(\tau) = \frac{1}{N}\sum_{i=1}^{N}[(x_i - \langle x\rangle)^2 + (y_i - \langle y\rangle)^2], \qquad (1)$$

where $N$ is the total number of time steps in each trajectory, $x_i$ and $y_i$ are the projection of the position of each trajectory step on the $x-$ and $y-$axis, respectively, and $\langle x\rangle$ and $\langle y\rangle$ are their mean values. Physically, $R_g$ quantifies the size of a GEMs trajectory generated during the time lapse $\tau$. A cutoff value of $(R_g')_c = 0.3$ was used in the experiment, below which the GEMs trajectories are treated as immobile ones [7]. Here $R_g' = R_g/\langle R_g\rangle$ is the normalized radius of gyration, with $\langle R_g\rangle$ being the mean value of $R_g$. Mean squared displacements (MSDs), $\langle\Delta\mathbf{r}^2(\tau)\rangle = \langle(\mathbf{r}(t+\tau) - \mathbf{r}(t))^2\rangle$, were then computed from the mobile trajectories and fitted to $\langle\Delta\mathbf{r}^2(\tau)\rangle = 4D\tau$ to obtain the diffusion coefficients of GEMs $D$ in BEAS-2B cells.

To characterize the more complicated motions of Rab6A-positive secretory vesicles and Rab5-positive early endosomes, we used a method that could automatically identify and extract the states of diffusive "jiggling" movement and the states of directed "runs", as described previously [10]. Briefly, we determined the motion state of an arbitrary point in the trajectory by analyzing the turning angles around it at different time scales. Directed runs longer than 20 time-steps (2 s) were used to compute the run velocity. Trajectories containing no directed "runs" were considered as immobile. The mobile ratio was defined as the ratio of the number of mobile trajectories over the total number of trajectories in a single cell. The mobile ratio and run time/total travel time ratio generally characterize how active the motors are in moving vesicles from the "pauses" to the "runs" states at the whole cell level.

***Ex vivo* acetylation assay.** *Ex vivo* acetylation was performed according to a previously published protocol [11]. BEAS-2B cells were first washed with PEM buffer (80mM PIPES, 1mM EGTA, 1mM $MgCl_2$, pH 6.8), and then incubated in PEM buffer supplemented with 1% Triton X-100 and 10 $\mu$M taxol for 1 min to extract the cell cytosol while maintaining the microtubules. Cells were washed again with PEM buffer supplemented with 10 $\mu$M taxol. Next, cells were incubated in PEM buffer supplemented with 10 $\mu$M taxol, 100 nM acetyl coenzyme A

(Sigma) and the indicated concentration of recombinant $\alpha$TAT1 at 37°C for 8 min for acetylation. This reaction was stopped by fixation in −20 °C methanol for 10 min and the cells were then used for immunostaining to determine their acetylation level.

**Statistics.** Data are expressed as mean ± s.e.m. unless specified otherwise. Graphs were created using Origin. Statistical tests were performed with two-tailed unpaired Student's t-test. The statistical details of each experiment can be found in the figure legends.

#### II. SUPPLEMENTARY FIGURES

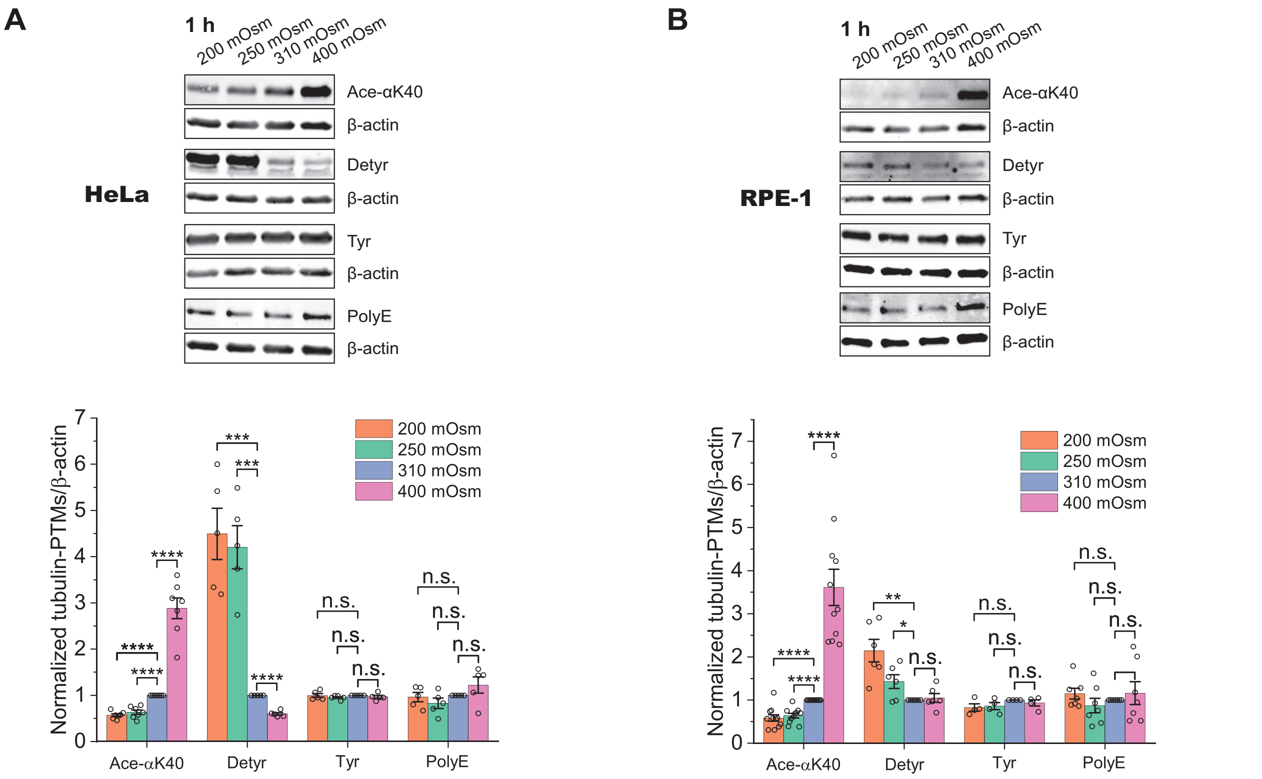


FIG. S1. **Cytoplasm crowdedness differentially tunes tubulin post-translational modifications in HeLa and RPE-1 cells.** Western-blots (WB) and corresponding quantification showing relative changes of tubulin PTMs in response to different osmotic treatments for 1 h in HeLa cells (A) and RPE-1 cells (B). Acetylation at Lys40 (Ace-$\alpha$K40), detyrosination (Detyr), tyrosination (Tyr), and polyglutamylation (PolyE) were assessed relative to the loading control, $\beta$-actin. For HeLa cells, Ace-$\alpha$K40, n = 7, ****$p < 0.0001$; Detyr, n = 5, ***$p = 0.00023$ and 0.00013, ****$p < 0.0001$; Tyr, n = 5; PolyE, n = 5. n.s. indicates $p > 0.05$. For RPE-1cells, Ace-$\alpha$K40, n = 11, ****$p < 0.0001$; Detyr, n = 6, **$p = 0.0014$, *$p = 0.022$; Tyr, n = 4; PolyE, n = 7. n.s. indicates $p > 0.05$. Error bars indicate s.e.m. All $p$-values were calculated using a student's t-test.

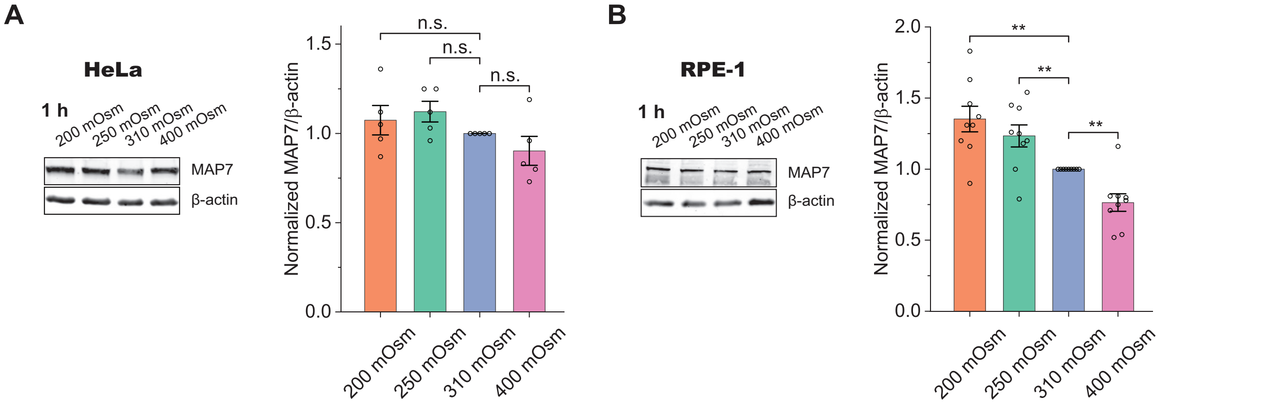


FIG. S2. **Hypoosmotic shifts upregulate MAP7 and hyperosmotic shifts downregulate MAP7 in RPE-1, but not in HeLa cells.** Western-blots and corresponding quantification showing the relative changes of MAP7 in response to osmotic treatments for 1 h in HeLa (A) and RPE-1 (B) cells. $\beta$-actin, loading control. HeLa: n = 5, no significant changes; RPE-1: n = 9, **$p = 0.0012$, 0.0085 and 0.0015. Error bars indicate s.e.m. All $p$-values were calculated using a student's t-test.

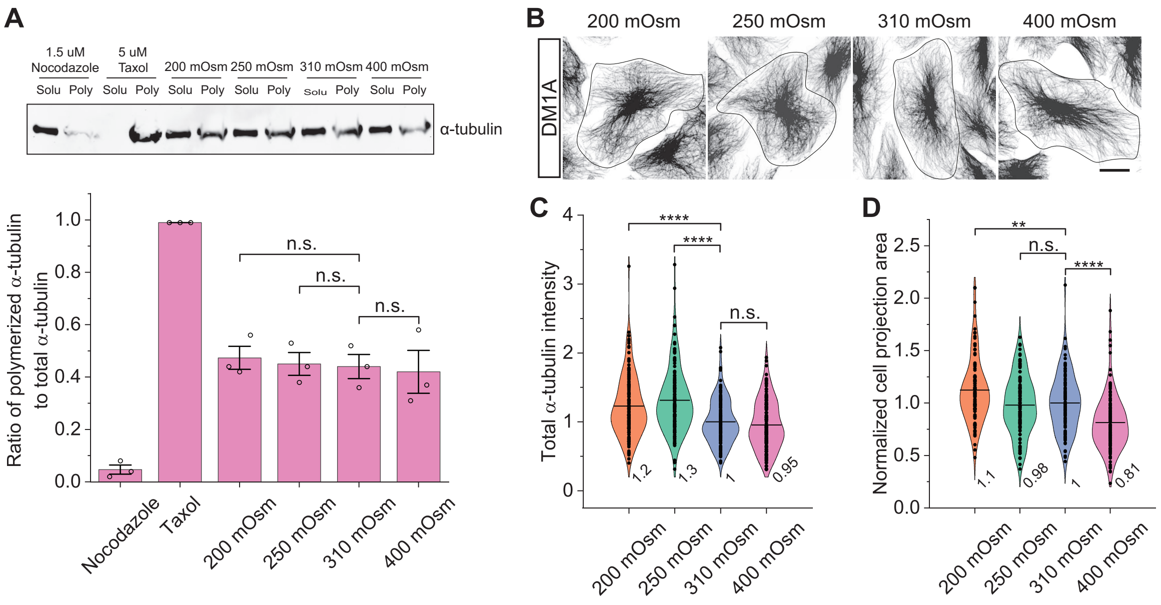


FIG. S3. **Modulation of cytoplasm crowdedness does not alter polymerized $\alpha$-tubulin in BEAS-2B cells in the experimental conditions.** (A) Western-blots (WB) and corresponding quantification showing the changes of the ratio of polymerized $\alpha$-tubulin to total $\alpha$-tubulin in response to nocodazole (1.5 $\mu$M), taxol (5 $\mu$M), and different osmotic treatments for 1 hr in BEAS-2B cells. Soluble (Solu) and polymerized (Poly) $\alpha$-tubulin were fractionated based on a previously published method [2]. n = 3, n.s. indicates $p > 0.05$. Error bars indicate s.e.m. (B-D) Representative images and quantification comparing the relative fluorescence intensity of $\alpha$-tubulin (C) and cell projection area (D) for cells treated with different osmotic solutions for 1 hr. n = 139, 141, 143 and 139 cells from 3 independent experiments. Cell projection area: $**p = 0.0030$, $****p < 0.0001$; total $\alpha$-tubulin intensity: $****p < 0.0001$. Scale bars: 20 $\mu$m. All graphs display all data points with means. All $p$-values were calculated using a student's t-test.

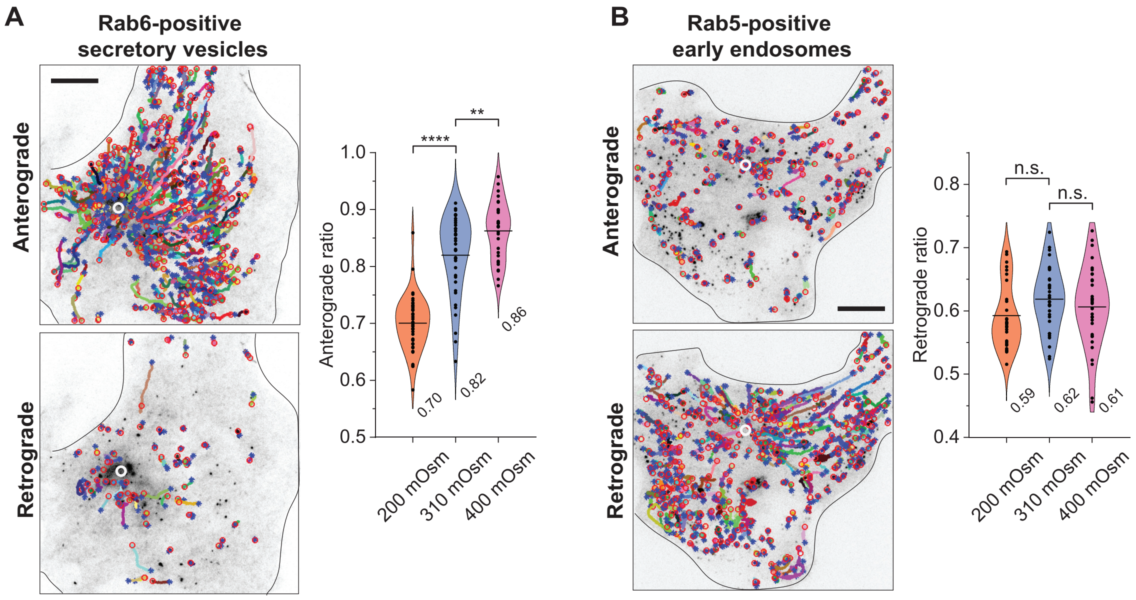


FIG. S4. **Cytoplasm crowdedness tunes transport direction of Rab6-positive secretory vesicles.** (A) Representative trajectories (colored lines) of EGFP-Rab6A-positive secretory vesicles in a BEAS-2B cell (10 fps for 3 min) showing the anterograde and retrograde fractions of all mobile movements observed (left) and quantification of the anterograde transport ratio in response to different osmotic treatments (right). $^{**}p = 0.0065$, $^{****}p < 0.0001$. The total trajectories analyzed from the mobile fraction are n = 29688, 32248 and 22572 from n = 42, 40 and 34 cells from 3 independent experiments. Each color represents an individual trajectory. Blue stars and red circles mark the start and end point of each trajectory, respectively. White circles indicate microtubule organizing center. Scale bar: 10 $\mu$m. (B) Representative trajectories (colored lines) of EGFP-Rab5-positive early endosomes in a BEAS-2B cell (10 fps for 3 min) showing the anterograde and retrograde fractions of all mobile movements observed (left) and quantification of the retrograde transport ratio in response to different osmotic treatments (right). The total trajectories analyzed from the mobile fraction are n = 33130, 28305 and 25248 from n = 37, 34 and 36 cells from 3 independent experiments. All graphs display all data points with means. All $p$-values were calculated using a student's t-test.
